# Transient tubular membrane injury is converted into persistent fibrosis through a macrophage–fibroblast relay

**DOI:** 10.64898/2026.09.28.755241

**Authors:** Peihan Wang, Yuan Ruan, Ting Rao, Bojun Li, Fan Cheng, Wei Li

**Author notes:** Correspondence: Fan Cheng, MD, PhD, Professor, Department of Urology, Renmin Hospital of Wuhan University, Wuhan, Hubei 430060, P.R. China. Wei Li, MD, PhD, Professor, Department of Anesthesiology, Renmin Hospital of Wuhan University, Wuhan, Hubei 430060, P.R. China. Peihan Wang and Yuan Ruan contributed equally to this work.

## Abstract

A central unresolved problem in acute kidney injury (AKI) is how a transient epithelial insult is converted into persistent fibrotic repair that drives progression toward chronic kidney disease. Although tubular, immune and stromal responses are individually well recognized, the mechanistically continuous relay linking these three compartments has remained unresolved. Here we define a directional information-conversion pathway spanning injured tubular epithelial cells, macrophages and fibroblasts. Time-resolved single-cell profiling identified a TREM2/SPP1 macrophage state that emerged as post-ischemic repair became fibrotic. Tubular injury remodeled phosphatidylserine (PS) species, increased oxidized PS and PS externalization, and generated a membrane-lipid context for TREM2 sensing, supported by PS masking, defined liposomes and orthogonal binding assays. Structure–function analysis separated R47-dependent phospholipid recognition from K183-dependent DAP12 coupling, linking membrane damage to SYK activation and NF-κB/AP-1–driven Spp1 transcription. Macrophage-derived SPP1 then activated ITGAV/ITGB1-associated FAK–SMAD2/3 signaling, fibroblast contraction and matrix production. Crucially, repair-phase SPP1 neutralization and post-injury Cx3cr1-lineage Trem2 deletion attenuated fibrosis, with delayed Trem2 deletion protective in both ischemic and obstructive injury. Human single-cell, spatial, urinary and tissue analyses showed cross-species and cross-scale convergence. These findings resolve how transient tubular membrane damage is successively transformed across three cellular compartments into persistent, yet interruptible, fibrotic repair.

## Introduction

Fibrosis can persist long after the epithelial insult that initiated it has subsided. This temporal disconnect is central to the transition from acute kidney injury (AKI) to chronic kidney disease (CKD): a transient tubular lesion must somehow be converted into a durable multicellular program of maladaptive repair. Kidney regeneration and fibrosis emerge from coordinated interactions among tubular epithelial cells, immune populations, and stromal cells. Persistently injured tubules acquire inflammatory and profibrotic states, sustained immune activation reshapes tissue macrophages, and activated fibroblasts deposit extracellular matrix[1–3]. Spatially organized injury niches further show that these responses are physically integrated within diseased kidney tissue[3]. Yet these compartments are still commonly viewed as parallel consequences of injury or as isolated pairwise interactions. What remains unresolved is the information flow that connects them: how membrane damage in an injured tubule is encoded, how that signal is decoded and transformed by macrophages, and how the resulting immune output is retransmitted to fibroblasts to sustain fibrosis after the initiating insult has passed.

Macrophages are uniquely positioned to perform this intermediate conversion because their state is continuously shaped by local damage, metabolic stress, and membrane-derived signals. Across fibrotic organs, recurrent macrophage populations are enriched for SPP1, GPNMB, TREM2, LGALS3, and lipid-handling genes[4]. SPP1-expressing macrophages can promote myofibroblast activation through macrophage–stromal communication, while Spp1+Fn1+Arg1+ macrophages accumulate in injured kidneys and their depletion attenuates fibrosis[5, 6]. Related TREM2/SPP1-rich and lipid-associated macrophage programs have also been identified in chronic kidney disease and other inflammatory renal states[7, 8]. These observations establish macrophages as a plausible bridge between parenchymal injury and stromal remodeling, but they leave the critical mechanistic steps unresolved: the upstream damage signal, the receptor logic that converts this signal into a new macrophage state, and the stromal pathway that executes the resulting fibrotic output. Triggering receptor expressed on myeloid cells 2 (TREM2) provides a candidate molecular interface between membrane injury and macrophage-state conversion. TREM2 is a lipid-responsive immunoreceptor that signals through DAP12 and SYK, recognizes anionic membrane lipids including phosphatidylserine (PS), and contains a positively charged ligand-interacting surface centered on Arg47[9, 10]. PS is normally confined to the inner leaflet of the plasma membrane but becomes externally accessible during cellular stress and injury, including ischemia–reperfusion contexts[11, 12]. Yet the biological consequences of TREM2 signaling in the kidney are strikingly context dependent. TREM2 can support macrophage survival and injury resolution in acute settings, whereas TREM2-expressing macrophage programs have also been associated with chronic remodeling and the AKI-to-CKD transition[13, 14]. This apparent paradox raises a temporal possibility: damage sensing that is adaptive during acute injury may become pathogenic when membrane-derived signals and macrophage activation persist into the repair phase.

Here we show that tubular membrane injury is not simply followed by fibrosis, but is progressively converted into it through a directional macrophage–fibroblast relay. Time-resolved single-cell analysis identifies a TREM2/SPP1 macrophage state that emerges as post-ischemic repair becomes fibrotic. Tubular injury remodels PS/OxPS species and increases PS externalization, creating a membrane-damage context decoded by TREM2 through R47-dependent phospholipid recognition and K183-dependent DAP12 coupling. This signal is then transcriptionally converted through DAP12–SYK–NF-κB/AP-1 into Spp1 expression, and macrophage-derived SPP1 is retransmitted to kidney fibroblasts through ITGAV/ITGB1-associated FAK–SMAD2/3 signaling. Crucially, SPP1 neutralization and Trem2 deletion initiated only after kidney injury still suppress subsequent fibrosis, with delayed Trem2 deletion protective across both ischemic and obstructive injury. Human single-cell, spatial, urinary, and tissue analyses provide cross-species and cross-scale support for the same macrophage–stromal architecture. Together, these findings define a continuous information-conversion process—from epithelial damage encoding, through macrophage decoding and transcriptional transformation, to stromal execution—and show that this process remains biologically active and interruptible after injury.

## Methods

### Study design and animal models

Male C57BL/6 mice underwent unilateral renal ischemia–reperfusion injury (IRI; 45-min left renal-artery occlusion with the contralateral kidney retained) or unilateral ureteral obstruction (UUO). IRI kidneys were collected at 1 and 28 d and UUO kidneys at 14 d. For repair-phase Trem2 deletion, Trem2fl/fl mice (Jackson Laboratory, stock 029853) were crossed with Cx3cr1-CreERT2 mice (stock 021160). Tamoxifen was administered on days 3–7 after IRI or days 1–5 after UUO; Cre-negative Trem2fl/fl littermates received the same regimen. Trem2 deletion was verified in sorted CD45□ CD11b□ F4/80□ kidney macrophages. In a separate IRI cohort, SPP1-neutralizing antibody or isotype IgG was administered intraperitoneally at 10 mg/kg every 3 d from days 3–27. Animal experiments were approved by the Animal Welfare and Ethics Review Committee of Renmin Hospital of Wuhan University (WDRM20200308A).

### Single-cell transcriptomics and cell–cell communication

Kidneys from Sham, IRI-1 d, and IRI-28 d mice (n = 3 per group) underwent single-cell RNA sequencing to define the temporal emergence of macrophage states across acute injury and chronic repair. After quality control and Seurat-based integration[15], macrophages were reclustered into C1–C13. The C3 program was quantified with AUCell using the predefined macrophage signature[16], and CellChat was used to infer SPP1 ligand–receptor communication between macrophages and stromal cells[17].

### Cell models, tubular injury, and lipid analyses

TCMK-1 cells and primary mouse kidney fibroblasts were obtained from Wuhan Pricella Biotechnology. Bone marrow-derived macrophages (BMDMs) were differentiated from C57BL/6 marrow for 7 d with 25 ng/ml M-CSF. TCMK-1 cells underwent 24 h hypoxia (1% O□) followed by 24 h reoxygenation, with ferrostatin-1 (1 μM) where indicated. Tubular lipid injury was assessed by LCN2/ACSL4/GPX4 immunoblotting, 4-HNE immunofluorescence, C11-BODIPY imaging, Annexin V-FITC/PI flow cytometry, and untargeted lipidomics (n = 5 per group) using high-resolution LC–MS/MS. Tubular debris was generated by three freeze–thaw cycles, with exposed PS masked by Annexin V where indicated. Native PS was obtained from Avanti Polar Lipids, OxPS generated by Fenton oxidation, and PC-, PS-, and OxPS-containing liposomes prepared for stimulation.

### TREM2 structure–function and signaling assays

BMDMs were transfected with Trem2- or Tyrobp/Dap12-targeting siRNAs. Trem2-depleted cells were rescued with wild-type TREM2, the extracellular R47A ligand-sensing mutant, or the transmembrane K183A DAP12-coupling mutant. Membrane fractionation, reciprocal co-immunoprecipitation, liposome co-sedimentation, and lipid–protein overlay assays assessed receptor localization, TREM2–DAP12 association, and lipid binding. R406, BAY 11-7082, and SP600125 inhibited SYK, NF-κB, and JNK/AP-1 signaling, respectively. Spp1 transcription was assessed by p65/c-Jun ChIP-qPCR and wild-type or binding-site-mutant promoter luciferase reporters; secreted SPP1 was measured by ELISA.

### Macrophage–fibroblast experiments and tissue analyses

BMDM-conditioned media were collected for 24 h and applied to primary mouse kidney fibroblasts for 24 h. SPP1-neutralizing antibody, recombinant mouse SPP1 (100 ng/ml), ITGAV/ITGB1 blocking antibodies (10 μg/ml), and Itgav/Itgb1 siRNAs were used to interrogate fibroblast responses. FAK–SMAD2/3 signaling and α-SMA, COL1A1, and FN1 were assessed by immunoblotting or immunofluorescence, and collagen-gel contraction followed for up to 7 d. Kidney fibrosis was quantified by Masson’s trichrome, Sirius Red, α-SMA immunofluorescence, and hydroxyproline.

### Human datasets and clinical samples

Human kidney single-cell and spatial transcriptomic data were obtained from GSE211785[18]. Monocyte/macrophage populations were reclustered to identify TREM2-high C4 and murine C3-like C5 states; TREM2□SPP1□ cells were defined from raw counts and tested for enrichment by donor-stratified Mantel–Haenszel analysis. Spatial analyses retained the original Mac and Fib/MyoFib annotations and examined macrophage SPP1/GPNMB and stromal ITGAV/ITGB1/matrix programs. The clinical cohort included 8 healthy male controls and 20 male patients with CKD; urinary-cell TREM2, SPP1, and CD68 were measured by qPCR. Kidney tissue included 6 non-neoplastic reference and 8 CKD samples, with matched urine–tissue analyses in CKD. The human study was approved by the Ethics Committee of Renmin Hospital of Wuhan University (WDRY2021-KS047).

### Statistics

GraphPad Prism 10.1.2 (GraphPad Software, USA) was used for statistical analysis. Student’s t test (unpaired, two-tailed) was used for prespecified comparisons between two independent experimental groups unless otherwise specified. When experiments contained multiple treatment conditions, statistical testing was restricted to prespecified biologically relevant pairwise comparisons, as indicated in the corresponding figure legends, rather than exhaustive comparisons among all possible group pairs.

For human cohort analyses, clinical biochemical variables and histological measurements were compared between two independent groups using two-tailed Mann–Whitney U tests. For urinary qPCR analyses, statistical comparisons were performed on ΔCt values using two-tailed unpaired Student’s t tests, whereas relative expression values calculated using the 2^−ΔΔCt method are presented for visualization. Spearman’s rank correlation analysis was used to evaluate associations between molecular, histological, and clinical variables. Donor-stratified enrichment of TREM2□SPP1□ macrophages in the human single-cell RNA-sequencing dataset was assessed using the Mantel–Haenszel method, with donor identity treated as the stratification variable.

All statistical tests were two-tailed, with P < 0.05 considered statistically significant. Independent biological replicates, rather than technical replicates or individual microscopic fields, were treated as the experimental unit. Cell-based experiments were independently repeated at least three times, and sample sizes for animal and human studies are provided in the corresponding figure legends.

## Results

### A TREM2/SPP1 macrophage state progressively emerges as repair becomes maladaptive

We first asked whether the macrophage program of interest was a static injury feature or a state that emerged during maladaptive repair. Single-cell RNA sequencing of Sham, IRI-1 d, and IRI-28 d kidneys resolved 13 macrophage subclusters (Fig. 1A–D). C3 showed the strongest chronic-stage enrichment, increasing from 2.2% of macrophages in Sham to 8.7% at IRI-1 d and 25.1% at IRI-28 d; 85.1% of C3 cells originated from IRI-28 d kidneys (Fig. 1E). Focused analysis further showed C3 co-expression of Lgals3, Atp6v0d2, Ctsl, Syngr1, Trem2, Spp1, Fabp5, Ctsd, Mfge8, and Gpnmb, with a signature score that increased progressively across injury stages (Fig. 1F,G). Thus, C3 represents a TREM2/SPP1-rich macrophage state increasingly enriched during maladaptive repair.

**Figure 1.**
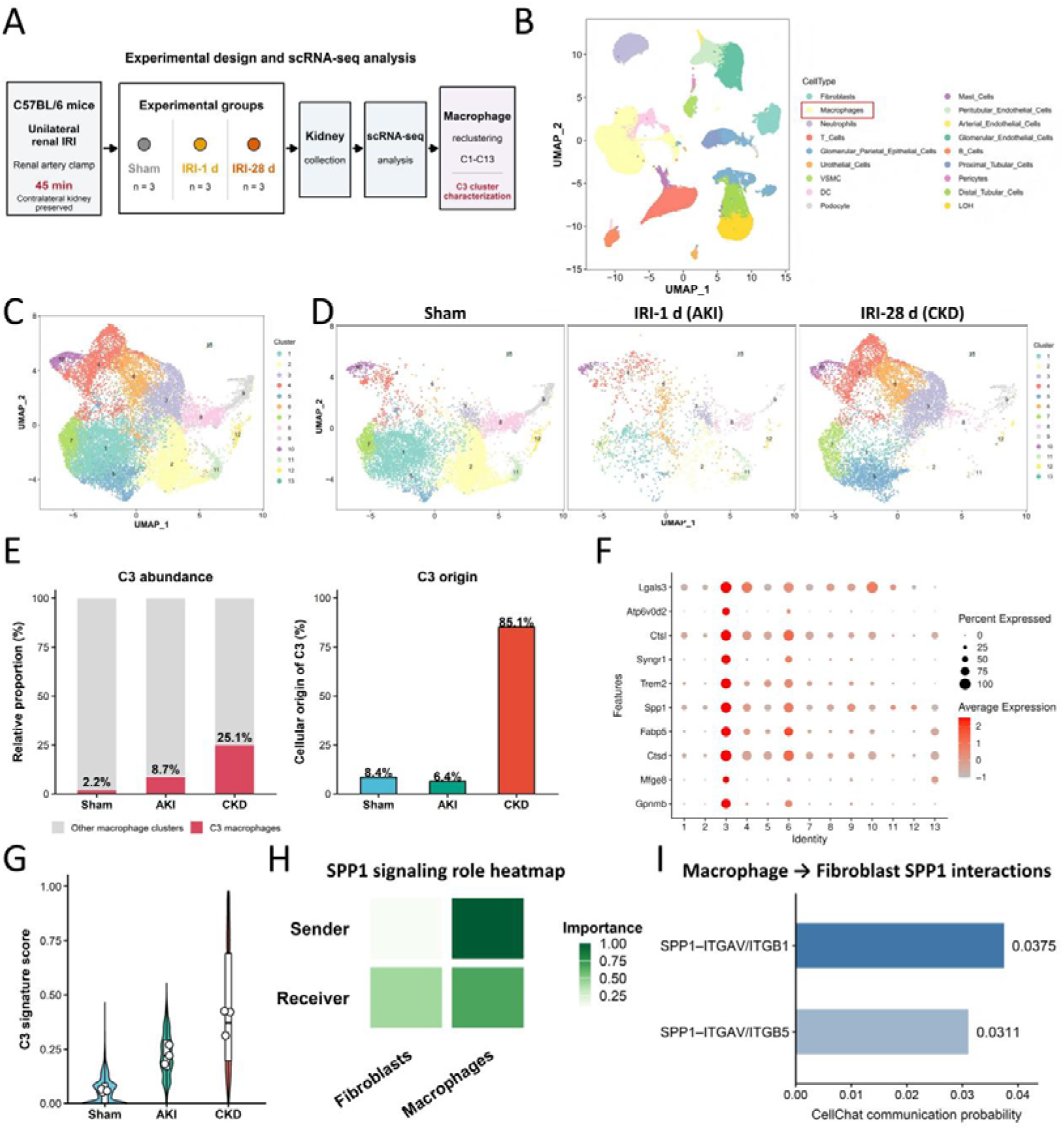
A TREM2/SPP1 macrophage state progressively emerges as repair becomes maladaptive. (A) Schematic overview of the experimental design and single-cell RNA sequencing workflow. C57BL/6 mice underwent unilateral renal ischemia–reperfusion injury (IRI) by renal artery clamping for 45 min with the contralateral kidney preserved. Kidneys were collected from Sham, IRI-1 d (AKI), and IRI-28 d (CKD) groups (n = 3 per group) for scRNA-seq analysis, followed by macrophage reclustering into 13 subclusters (C1–C13). (B) UMAP visualization of major renal cell populations identified in the integrated single-cell dataset. Macrophages are indicated within the global cellular landscape. (C) UMAP visualization of reclustered macrophages, resolving 13 transcriptionally distinct subclusters (C1–C13). (D) Group-separated UMAP plots showing the distribution of macrophage subclusters in Sham, IRI-1 d, and IRI-28 d kidneys. (E) Left, relative abundance of C3 macrophages among total macrophages in Sham, AKI, and CKD kidneys. C3 macrophages accounted for 2.2%, 8.7%, and 25.1% of macrophages, respectively. Right, cellular origin of the C3 population across experimental groups, showing that 85.1% of C3 macrophages were derived from CKD kidneys, compared with 8.4% from Sham and 6.4% from AKI kidneys. (F) Dot plot showing expression of representative C3-associated genes, including Lgals3, Atp6v0d2, Ctsl, Syngr1, Trem2, Spp1, Fabp5, Ctsd, Mfge8, and Gpnmb, across macrophage subclusters. Dot size denotes the proportion of expressing cells and color intensity denotes mean expression. (G) Distribution of the C3 transcriptional signature score across Sham, AKI, and CKD macrophages, showing progressive enrichment of the C3-associated program during the AKI-to-CKD transition. (H) CellChat-derived SPP1 signaling role heatmap showing the relative sender and receiver activities of macrophages and fibroblasts within the SPP1 signaling network. (I) Predicted macrophage-to-fibroblast SPP1 ligand–receptor interactions identified by CellChat. SPP1–ITGAV/ITGB1 and SPP1–ITGAV/ITGB5 displayed communication probabilities of 0.0375 and 0.0311, respectively.

CellChat analysis identified a potential route linking this state to the stromal compartment. Macrophages showed prominent outgoing SPP1 signaling, whereas fibroblasts displayed a receiver-dominant profile, with predicted SPP1–ITGAV/ITGB1 and SPP1–ITGAV/ITGB5 interactions (Fig. 1H,I). Together, the discovery dataset links progressive emergence of the TREM2/SPP1-rich macrophage state with an SPP1-centered macrophage-to-fibroblast communication axis.

### The TREM2/SPP1 state localizes to the fibrotic phase of post-ischemic repair

We next anchored the temporal single-cell trajectory in kidney tissue. IRI-1 d kidneys showed acute tubular injury, whereas IRI-28 d kidneys displayed structural remodeling, increased Masson’s trichrome and Sirius Red staining, expanded interstitial α-SMA, and higher α-SMA, COL1A1, and FN1 abundance (Fig. 2A–D). Multicolor immunofluorescence showed increasing TREM2 and SPP1 signals within F4/80-rich regions, with greatest overlap at IRI-28 d (Fig. 2E), linking the transcriptional state to established fibrotic remodeling.

**Figure 2.**
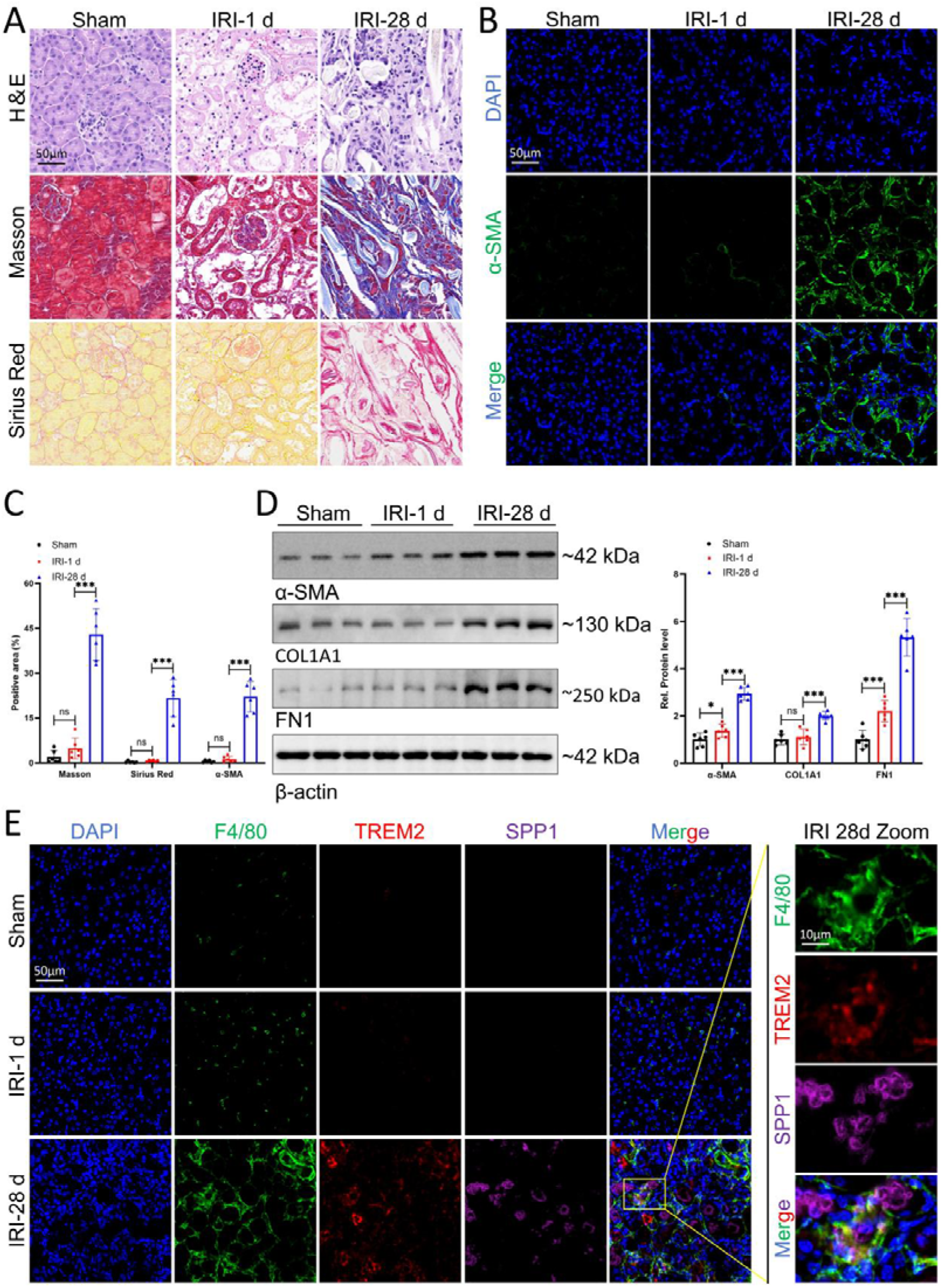
The TREM2/SPP1 macrophage state accumulates in the fibrotic phase of post-ischemic kidney repair. (A) Representative hematoxylin and eosin (H&E), Masson’s trichrome, and Sirius Red staining of kidney sections from Sham, IRI-1 d, and IRI-28 d mice. H&E staining illustrates acute tubular injury at 1 d and persistent structural remodeling at 28 d, whereas Masson’s trichrome and Sirius Red staining demonstrate prominent interstitial collagen deposition at the chronic stage. Scale bar, 50 μm. (B) Representative immunofluorescence images of α-SMA in kidneys from Sham, IRI-1 d, and IRI-28 d mice. Nuclei were counterstained with DAPI. Scale bar, 50 μm. (C) Quantification of Masson-positive area, Sirius Red-positive area, and α-SMA-positive area in the indicated groups. (D) Representative immunoblotting and densitometric quantification of α-SMA, COL1A1, and FN1 in kidney tissues from Sham, IRI-1 d, and IRI-28 d mice. β-actin was used as the loading control. Approximate molecular weights are indicated. (E) Representative multicolor immunofluorescence staining for F4/80 (green), TREM2 (red), and SPP1 (magenta) in kidneys from Sham, IRI-1 d, and IRI-28 d mice. Nuclei were counterstained with DAPI (blue). The enlarged view of the IRI-28 d kidney shows spatial overlap of TREM2 and SPP1 signals within F4/80-rich regions. Scale bars, 50 μm for overview images and 10 μm for the enlarged images. Data are presented as mean ± SD. Statistical significance between the indicated groups was assessed using a two-tailed unpaired Student’s t-test. ns, not significant; *P < 0.05; **P < 0.01, ***P < 0.001.

### Tubular injury encodes a phosphatidylserine-rich membrane damage signal

Having identified a macrophage state that progressively emerges during fibrotic repair, we next asked what upstream damage signal is presented by injured tubules to TREM2-expressing macrophages. In TCMK-1 cells, hypoxia/reoxygenation (H/R) induced membrane oxidative stress, with increased LCN2, altered ACSL4 and GPX4 abundance, and accumulation of 4-HNE and oxidized C11-BODIPY signals; ferrostatin-1 (Fer-1) partially attenuated these changes (Fig. 3A–D).

**Figure 3.**
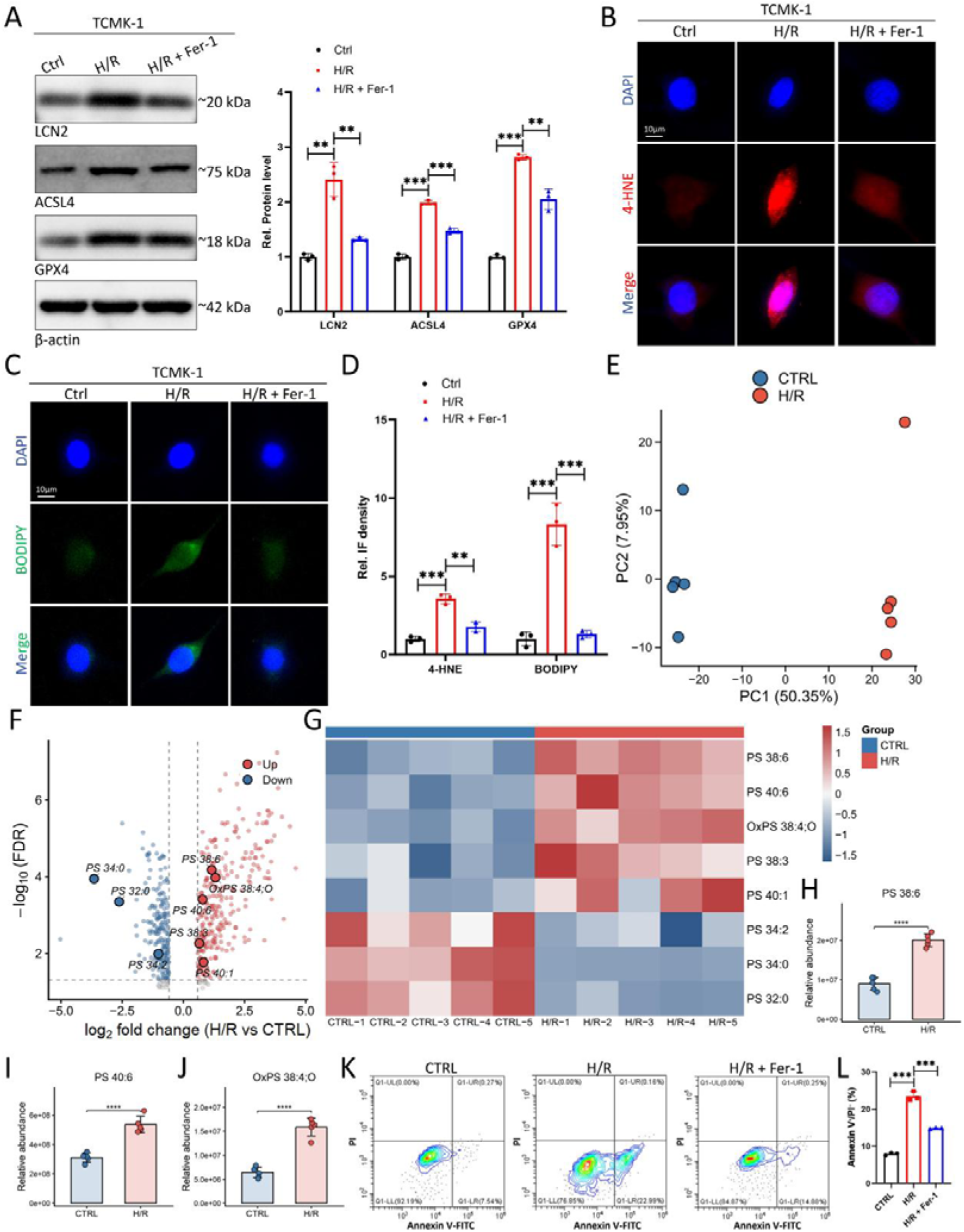
Tubular injury encodes a phosphatidylserine-rich membrane damage signal. (A) Representative immunoblots and densitometric quantification of LCN2, ACSL4, and GPX4 in TCMK-1 cells under control conditions, following hypoxia/reoxygenation (H/R), or following H/R in the presence of ferrostatin-1 (Fer-1). β-actin served as the loading control. Approximate molecular weights are indicated. (B) Representative immunofluorescence images of 4-hydroxynonenal (4-HNE) in TCMK-1 cells under the indicated conditions. Nuclei were counterstained with DAPI. Scale bar, 10 μm. (C) Representative C11-BODIPY fluorescence images showing H/R-induced lipid oxidative stress and its attenuation by Fer-1. Nuclei were counterstained with DAPI. Scale bar, 10 μm. (D) Quantification of relative 4-HNE and C11-BODIPY fluorescence intensity in the indicated groups. (E) Principal component analysis of untargeted lipidomic profiles from control and H/R-treated TCMK-1 cells (n = 5 biological replicates per group), showing separation between the two conditions. (F) Volcano plot of differential lipid species between H/R and control TCMK-1 cells. Selected phosphatidylserine (PS) and oxidized phosphatidylserine (OxPS) species are indicated. (G) Heatmap showing relative abundance patterns of selected PS and OxPS species across control and H/R samples. (H–J) Relative abundance of representative injury-associated phospholipid species, including PS 38:6 (H), PS 40:6 (I), and OxPS 38:4;O (J), in control and H/R-treated TCMK-1 cells. (K) Representative Annexin V-FITC/PI contour plots of TCMK-1 cells under control, H/R, and H/R + Fer-1 conditions. Annexin V□/PI□ cells were defined as cells with externalized phosphatidylserine while retaining plasma-membrane integrity. (L) Quantification of Annexin V□/PI□ cells in the indicated groups (n = 3 independent experiments). Data are shown as mean ± SD; indicated pairwise comparisons were performed using two-tailed unpaired Student’s t tests. **P < 0.01, ***P < 0.001, ****P < 0.0001.

Untargeted lipidomics showed clear separation between control and H/R-treated cells and broad lipid remodeling (Fig. 3E,F). Phosphatidylserine (PS)- and oxidized PS (OxPS)-related species were prominently altered (Fig. 3G), with PS 38:6, PS 40:6, and OxPS 38:4;O significantly increased after H/R (Fig. 3H–J).

H/R also increased cell-surface PS accessibility by Annexin V-FITC/PI flow cytometry. Annexin V□/PI□ cells increased from approximately 8% in controls to approximately 23% after H/R and were partially reduced to approximately 15% by Fer-1 (Fig. 3K,L). Because these cells retain plasma-membrane integrity while exposing PS on the outer leaflet, the data directly demonstrate increased extracellular PS accessibility during tubular oxidative injury.

Together, these findings define tubular injury as a membrane-damage state characterized by lipid oxidation, PS/OxPS remodeling, and PS externalization, providing a candidate phospholipid signal for lipid-responsive macrophages.

### TREM2 decodes injury-exposed phosphatidylserine into a macrophage SPP1 program

We next tested whether PS within injured tubular membranes contributes to macrophage activation and whether defined PS/OxPS is sufficient to reproduce this signal. H/R-derived tubular debris increased TREM2 abundance, TREM2–DAP12 signaling, SYK phosphorylation, and SPP1 in BMDMs compared with control debris (Fig. 4A–C). Suppressing tubular lipid oxidation with ferrostatin-1 or masking exposed PS with Annexin V attenuated these responses. Conversely, defined PS and OxPS liposomes increased p-SYK and SPP1, with OxPS producing the strongest response and PC showing little effect (Fig. 4D–F). Trem2 silencing blunted PS/OxPS-induced SYK activation and SPP1 without materially altering DAP12 abundance. Reciprocal co-immunoprecipitation further showed that H/R debris enhanced TREM2–DAP12 complex formation, which was reduced by ferrostatin-1 or PS masking (Fig. 4G,H). H/R debris also increased secreted SPP1, and both interventions attenuated this response (Fig. 4I). Together, these data identify PS/OxPS-containing injured membranes as a biologically active upstream signal decoded through macrophage TREM2–DAP12–SYK.

**Figure 4.**
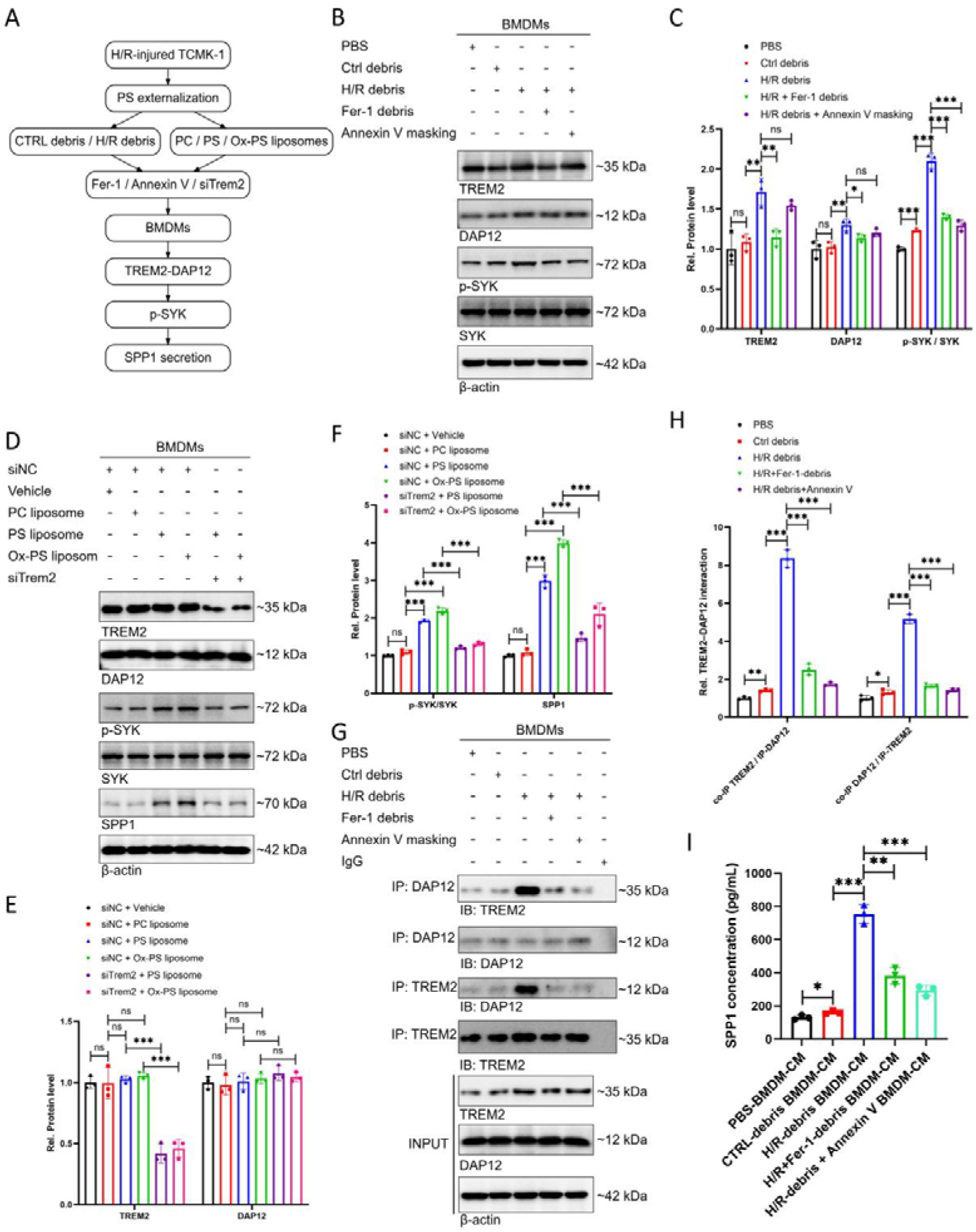
TREM2 decodes injury-exposed phosphatidylserine into DAP12–SYK activation and SPP1 production. (A) Schematic overview of the experimental strategy used to interrogate phosphatidylserine-dependent macrophage activation. Debris derived from control or H/R-injured TCMK-1 cells, as well as PC-, PS-, or oxidized PS (OxPS)-containing liposomes, were applied to bone marrow-derived macrophages (BMDMs). Ferrostatin-1 (Fer-1), Annexin V-mediated PS masking, and Trem2 silencing were used to assess the contribution of lipid oxidation, exposed PS, and TREM2, respectively. (B) Representative immunoblots of TREM2, DAP12, phosphorylated SYK (p-SYK), total SYK, and β-actin in BMDMs treated with PBS, control-cell debris, H/R-cell debris, debris derived from Fer-1-treated H/R cells, or H/R-cell debris subjected to Annexin V-mediated PS masking. (C) Densitometric quantification of TREM2, DAP12, and the p-SYK/SYK ratio under the conditions shown in (B). (D) Representative immunoblots of TREM2, DAP12, p-SYK, total SYK, SPP1, and β-actin in BMDMs exposed to vehicle, PC, PS, or OxPS liposomes in the presence of control siRNA (siNC), or to PS/OxPS liposomes following Trem2 silencing. (E) Densitometric quantification of TREM2 and DAP12 protein abundance under the conditions shown in (D). (F) Quantification of the p-SYK/SYK ratio and SPP1 protein abundance in liposome-treated BMDMs. (G) Reciprocal co-immunoprecipitation analysis of the TREM2–DAP12 complex in BMDMs treated with the indicated cell-debris preparations. DAP12 immunoprecipitates were immunoblotted for TREM2, and reciprocal TREM2 immunoprecipitates were immunoblotted for DAP12. Corresponding immunoprecipitation controls and input lysates are shown. (H) Quantification of TREM2–DAP12 association detected by reciprocal co-immunoprecipitation. (I) SPP1 concentrations in conditioned media from BMDMs treated with PBS, control-cell debris, H/R-cell debris, Fer-1-associated H/R-cell debris, or Annexin V-masked H/R-cell debris, as determined by ELISA. Data are shown as mean ± SD; indicated pairwise comparisons were performed using two-tailed unpaired Student’s t tests. ns, not significant; *P < 0.05, **P < 0.01, ***P < 0.001.

### R47 lipid recognition and K183 adaptor coupling define sequential steps in TREM2 damage decoding

We next resolved how TREM2 converts an extracellular membrane cue into intracellular signaling. After endogenous Trem2 depletion, rescue with wild-type TREM2, R47A, or K183A restored comparable total and membrane-associated receptor abundance (Fig. 5A–C), excluding major differences in expression or localization. In liposome co-sedimentation and lipid-overlay assays, wild-type TREM2-ECD-Fc preferentially bound PS and OxPS, whereas R47A markedly reduced both interactions (Fig. 5D,E).

**Figure 5.**
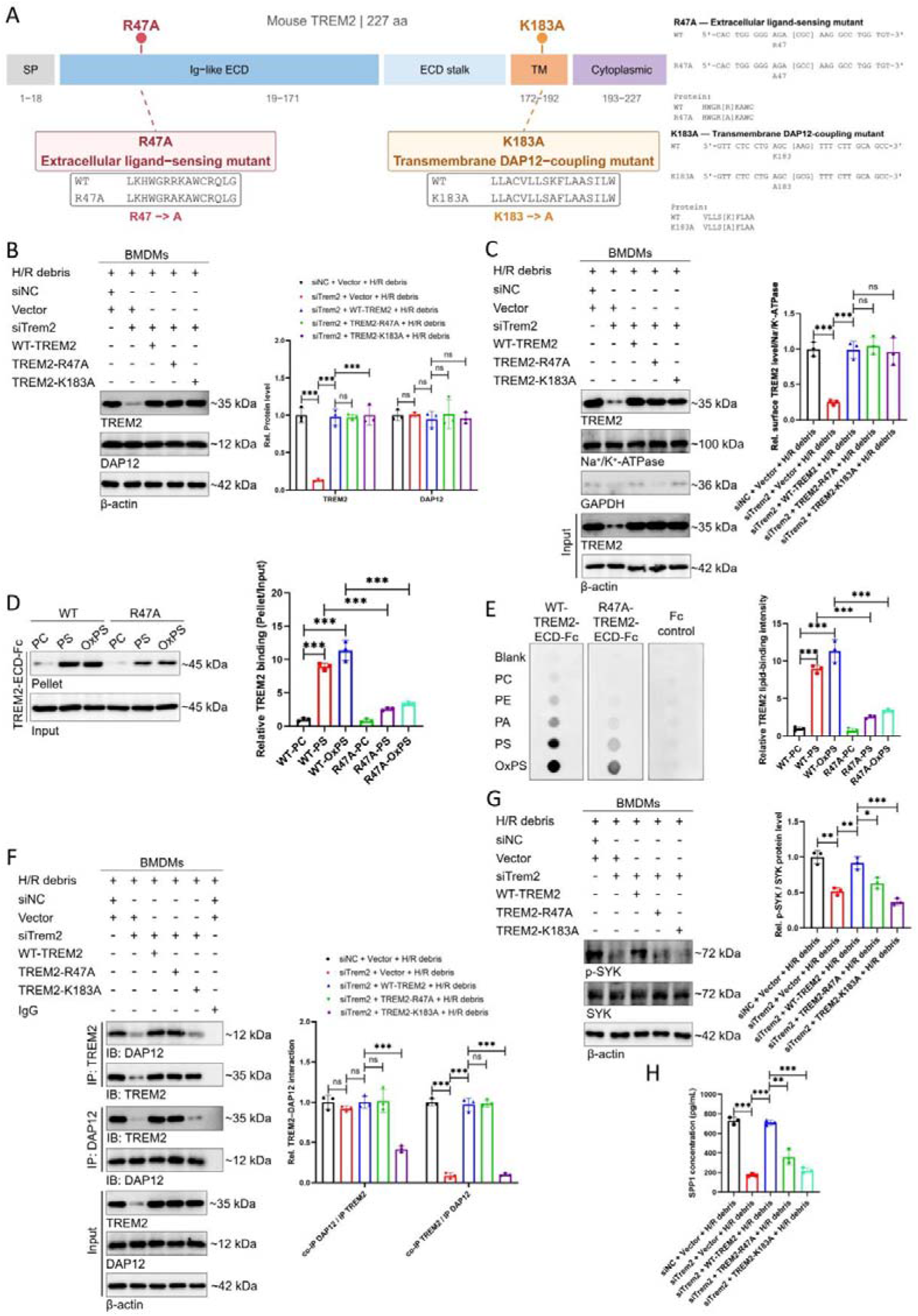
R47 lipid recognition and K183 DAP12 coupling define sequential steps in TREM2 damage decoding. (A) Schematic representation of mouse TREM2 domain organization and the engineered R47A and K183A mutations. R47A was introduced within the extracellular Ig-like domain as a ligand-sensing mutant, whereas K183A was introduced within the transmembrane region as a DAP12-coupling mutant. The corresponding nucleotide and amino-acid substitutions are shown. (B) BMDMs received control or Trem2-targeting siRNA, followed by rescue with empty vector, WT-TREM2, TREM2-R47A, or TREM2-K183A before stimulation with H/R-derived tubular debris. Representative immunoblots and densitometric quantification of TREM2 and DAP12 are shown. β-actin served as the loading control. (C) Membrane fractionation analysis of TREM2 membrane-associated abundance in BMDMs under the indicated conditions. Na+/K+-ATPase was used as a membrane-loading control and GAPDH as a cytosolic control. Total-cell input is shown below. Right, quantification of membrane-associated TREM2 normalized to Na+/K+-ATPase. (D) Liposome co-sedimentation assay assessing binding of recombinant WT- or R47A-TREM2-ECD-Fc to PC-, PS-, or OxPS-containing liposomes. Input and pellet fractions were analyzed for TREM2-ECD-Fc. Right, densitometric quantification expressed as relative TREM2 binding calculated from the pellet/input ratio. (E) Lipid–protein overlay assay assessing direct binding of WT-TREM2-ECD-Fc, R47A-TREM2-ECD-Fc, or Fc-only control to immobilized PC, PE, PA, PS, and OxPS. Representative blots from independent experiments are shown. Right, quantification of relative TREM2 lipid-binding intensity for the indicated PC, PS, and OxPS conditions. WT-TREM2 showed preferential binding to PS and OxPS, whereas R47A markedly reduced these interactions. (F) Reciprocal co-immunoprecipitation analysis of the TREM2–DAP12 complex in BMDMs following Trem2 knockdown and rescue with WT-TREM2, R47A, or K183A. TREM2 immunoprecipitates were immunoblotted for DAP12, and reciprocal DAP12 immunoprecipitates were immunoblotted for TREM2. IgG served as the negative immunoprecipitation control, and corresponding input lysates are shown. Right, densitometric quantification of reciprocal TREM2–DAP12 association. (G) Representative immunoblots and quantification of phosphorylated SYK and total SYK in BMDMs under the indicated rescue conditions. β-actin served as the loading control. (H) SPP1 concentrations in BMDM-conditioned media measured by ELISA following H/R debris stimulation under the indicated Trem2 knockdown and rescue conditions. Data are presented as mean ± SD from independent experiments. Statistical significance between the indicated groups was assessed using two-tailed Student’s t-tests. *P < 0.05, **P < 0.01, ***P < 0.001; ns, not significant.

Reciprocal co-immunoprecipitation showed that R47A largely preserved TREM2–DAP12 association, whereas K183A strongly disrupted it (Fig. 5F). Both mutants impaired H/R debris-induced SYK phosphorylation and SPP1 secretion despite preserved receptor abundance, with the strongest defect after K183A rescue (Fig. 5G,H). Thus, TREM2 damage decoding comprises two separable steps: R47-dependent phospholipid recognition followed by K183-dependent DAP12 coupling.

### TREM2–DAP12–SYK reprograms macrophage output through NF-κB/AP-1-dependent Spp1 transcription

Having defined ligand recognition and adaptor coupling, we next asked how TREM2 converts damage sensing into macrophage output. Trem2 or Dap12 silencing suppressed H/R debris-induced SYK phosphorylation and SPP1, placing the receptor–adaptor complex upstream of SPP1 induction (Fig. 6A–C). H/R debris increased p65 and c-Jun phosphorylation and nuclear accumulation, whereas SYK inhibition with R406 reduced both pathways and SPP1 (Fig. 6D). BAY 11-7082 and SP600125 preferentially suppressed p65 and c-Jun activation, respectively, and both reduced SPP1 protein and Spp1 mRNA (Fig. 6E,F).

**Figure 6.**
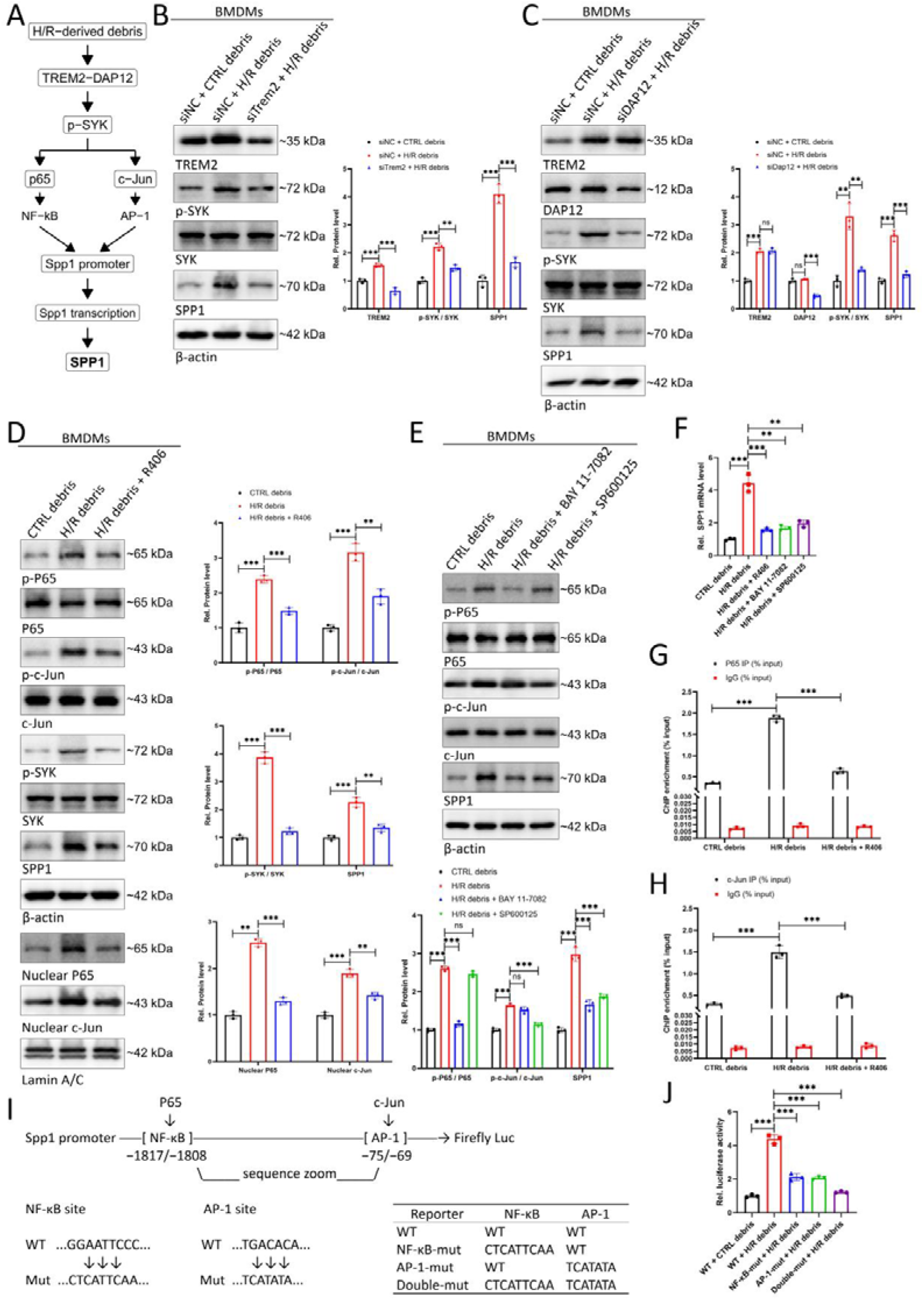
TREM2–DAP12–SYK reprograms macrophage output through NF-κB/AP-1-dependent Spp1 transcription. (A) Schematic representation of the proposed signaling cascade linking H/R-derived debris to TREM2–DAP12 activation, SYK phosphorylation, NF-κB/AP-1 activation, and subsequent Spp1 transcription in BMDMs. (B) Representative immunoblots and densitometric quantification of TREM2, phosphorylated SYK (p-SYK), total SYK, and SPP1 in BMDMs treated with control debris or H/R-derived debris in the presence of control siRNA (siNC) or Trem2 siRNA. β-actin served as the loading control. (C) Representative immunoblots and quantification of TREM2, DAP12, p-SYK, total SYK, and SPP1 in BMDMs following Dap12 silencing and stimulation with H/R-derived debris. (D) Effects of pharmacological SYK inhibition on downstream NF-κB and AP-1 signaling. BMDMs were exposed to control debris, H/R-derived debris, or H/R-derived debris in the presence of R406. Whole-cell lysates were analyzed for p-p65/p65, p-c-Jun/c-Jun, p-SYK/SYK, and SPP1, whereas nuclear fractions were analyzed for p65 and c-Jun accumulation. Lamin A/C was used as a nuclear loading control. Corresponding densitometric analyses are shown. (E) Representative immunoblots and quantification of p-p65/p65, p-c-Jun/c-Jun, and SPP1 in BMDMs treated with control debris, H/R-derived debris, H/R-derived debris plus the NF-κB inhibitor BAY 11-7082, or H/R-derived debris plus the JNK/AP-1 pathway inhibitor SP600125. (F) Relative Spp1 mRNA expression in BMDMs under the treatment conditions shown in (E). (G) ChIP-qPCR analysis of p65 occupancy at the Spp1 promoter in BMDMs treated with control debris, H/R-derived debris, or H/R-derived debris plus R406. IgG served as the negative immunoprecipitation control. Enrichment is expressed as percentage of input. (H) ChIP-qPCR analysis of c-Jun occupancy at the Spp1 promoter under the indicated conditions. (I) Schematic representation of the predicted NF-κB/p65 and AP-1/c-Jun response elements within the mouse Spp1 promoter and the corresponding wild-type and mutant luciferase reporter constructs. The NF-κB element is located at −1817/−1808 and the AP-1 element at −75/−69 relative to the transcriptional start site. Reporter constructs included wild-type, NF-κB-site mutant, AP-1-site mutant, and double-mutant promoters. (J) Relative luciferase activity of wild-type and mutant Spp1 promoter reporters following stimulation with control or H/R-derived debris. Data are shown as mean ± SD; indicated pairwise comparisons were performed using two-tailed unpaired Student’s t tests. ns, not significant; **P < 0.01; ***P < 0.001.

ChIP-qPCR showed increased p65 and c-Jun occupancy at the Spp1 promoter after H/R debris, which was reduced by R406 (Fig. 6G,H). Mutation of either the NF-κB or AP-1 response element diminished H/R debris-induced Spp1 promoter activity, with combined mutation producing the greatest suppression (Fig. 6I,J). Together, these data establish that TREM2–DAP12–SYK reprograms macrophage output through NF-κB/AP-1-dependent Spp1 transcription, converting a membrane-lipid signal into a soluble effector.

### SPP1 retransmits the decoded macrophage signal to fibroblasts through ITGAV/ITGB1–FAK–SMAD2/3

We next tested whether SPP1 transmits macrophage damage decoding to the stromal compartment. H/R debris increased SPP1 release from BMDMs, and conditioned medium from these macrophages activated primary mouse kidney fibroblasts, increasing FAK and SMAD2/3 phosphorylation together with COL1A1, FN1, and α-SMA (Fig. 7A–C). SPP1 neutralization reduced these responses and decreased α-SMA and fibronectin fluorescence (Fig. 7D,E).

**Figure 7.**
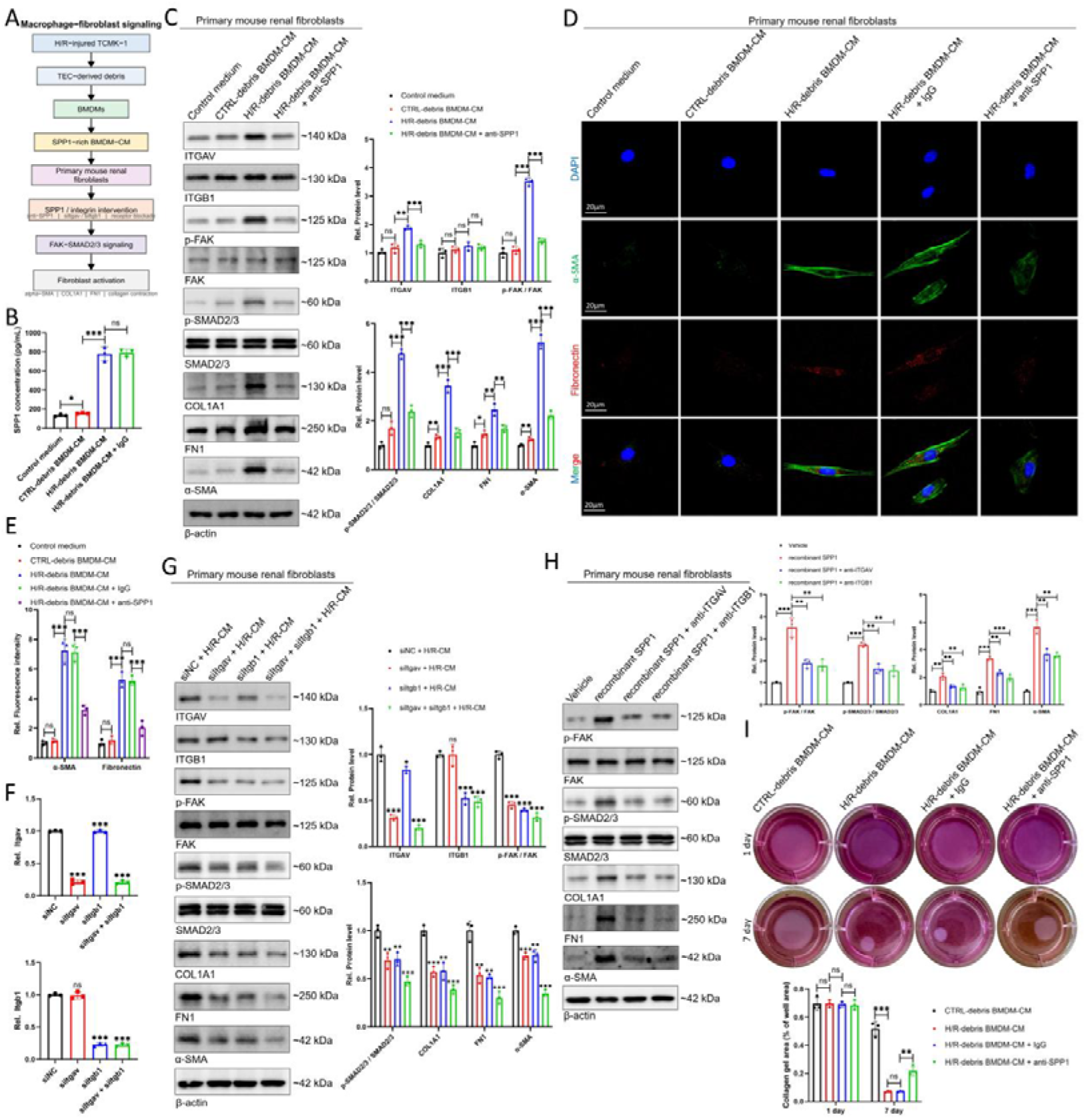
SPP1 retransmits the macrophage-decoded injury signal to kidney fibroblasts through ITGAV/ITGB1–FAK–SMAD2/3. (A) Schematic overview of the macrophage–fibroblast communication experiments. BMDMs were stimulated with debris derived from H/R-injured TCMK-1 cells, and the resulting conditioned medium (BMDM-CM) was applied to primary mouse kidney fibroblasts. SPP1 neutralization and genetic or antibody-mediated interference with ITGAV/ITGB1 were used to interrogate the downstream signaling pathway and fibroblast activation. (B) SPP1 concentrations in control medium and conditioned media from BMDMs exposed to control debris or H/R-derived debris, with IgG included as an antibody control where indicated, as determined by ELISA. (C) Representative immunoblots and densitometric quantification of ITGAV, ITGB1, phosphorylated FAK (p-FAK), total FAK, phosphorylated SMAD2/3 (p-SMAD2/3), total SMAD2/3, COL1A1, FN1, and α-SMA in primary mouse kidney fibroblasts treated with control medium, control-debris BMDM-CM, H/R-debris BMDM-CM, or H/R-debris BMDM-CM in the presence of an SPP1-neutralizing antibody. β-actin was used as the loading control. (D) Representative immunofluorescence images of α-SMA (green) and fibronectin (red) in primary mouse kidney fibroblasts subjected to the indicated conditioned-medium treatments, with IgG or SPP1-neutralizing antibody added where indicated. Nuclei were counterstained with DAPI (blue). Scale bar, 20 μm. (E) Quantification of α-SMA and fibronectin fluorescence intensity under the conditions shown in (D). (F) Validation of Itgav and Itgb1 silencing in primary mouse kidney fibroblasts following transfection with the indicated siRNAs. (G) Representative immunoblots and densitometric quantification of ITGAV, ITGB1, p-FAK/FAK, p-SMAD2/3/SMAD2/3, COL1A1, FN1, and α-SMA in primary mouse kidney fibroblasts treated with H/R-debris BMDM-CM following individual or combined silencing of Itgav and Itgb1. β-actin served as the loading control. (H) Representative immunoblots and quantitative analysis of FAK and SMAD2/3 activation and fibrotic protein expression in primary mouse kidney fibroblasts treated with recombinant SPP1 alone or in combination with blocking antibodies against ITGAV or ITGB1. (I) Representative images and quantification of collagen gel contraction by primary mouse kidney fibroblasts treated with control-debris BMDM-CM, H/R-debris BMDM-CM, H/R-debris BMDM-CM plus control IgG, or H/R-debris BMDM-CM plus SPP1-neutralizing antibody. Gel contraction was assessed at the indicated time points and expressed as collagen gel area relative to the well area. Data are shown as mean ± SD; indicated pairwise comparisons were performed using two-tailed unpaired Student’s t tests. ns, not significant; *P < 0.05, **P < 0.01, ***P < 0.001.

Depletion of Itgav or Itgb1, alone or together, similarly reduced conditioned-medium-induced FAK–SMAD2/3 signaling and fibrotic protein expression (Fig. 7F,G). Recombinant SPP1 activated the same pathway, whereas ITGAV or ITGB1 blockade attenuated the response (Fig. 7H). H/R-debris macrophage-conditioned medium also enhanced collagen-gel contraction, which was reversed by SPP1 neutralization (Fig. 7I). Together, these data establish SPP1–ITGAV/ITGB1–FAK–SMAD2/3 as the macrophage-to-fibroblast arm of the relay driving matrix remodeling.

### Repair-phase SPP1 neutralization interrupts stromal execution after AKI

We next asked whether the downstream relay remains important after injury. SPP1 neutralization began 3 d after IRI, with kidneys analyzed at day 28. IRI increased FAK and SMAD2/3 phosphorylation and α-SMA, COL1A1, and FN1, whereas anti-SPP1 reduced these responses (Fig. 8A). Masson’s trichrome, Sirius Red, interstitial α-SMA, and renal SPP1 staining were likewise reduced (Fig. 8B–H), together with kidney hydroxyproline, while the injured-to-contralateral kidney-weight ratio partially recovered (Fig. 8I,J). Thus, repair-phase SPP1 remains causally important to subsequent fibrosis.

**Figure 8.**
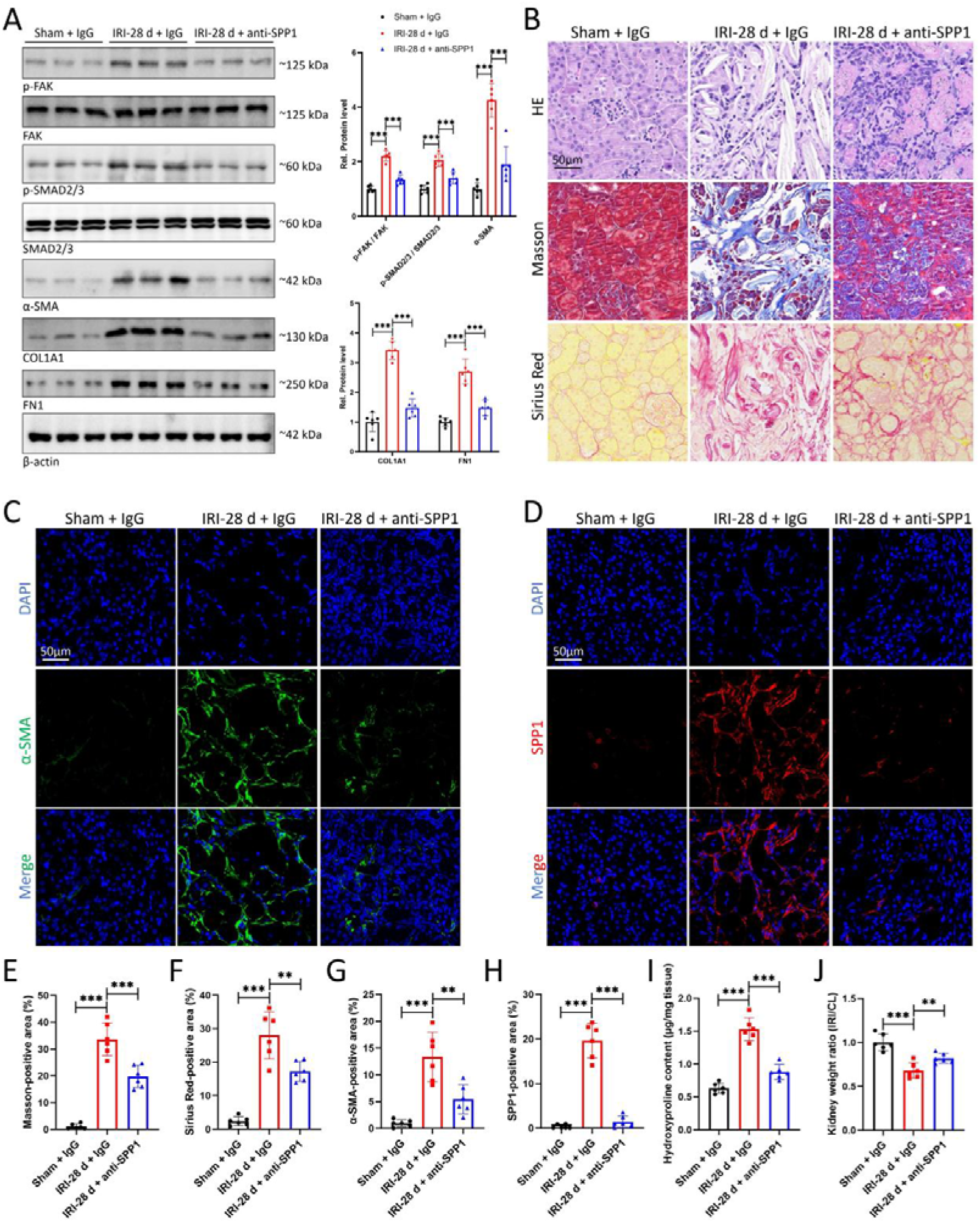
Repair-phase SPP1 neutralization interrupts stromal execution and attenuates kidney fibrosis. (A) Representative immunoblots and densitometric quantification of phosphorylated FAK (p-FAK), total FAK, phosphorylated SMAD2/3 (p-SMAD2/3), total SMAD2/3, α-SMA, COL1A1, and FN1 in kidneys from Sham + IgG, IRI-28 d + IgG, and IRI-28 d + anti-SPP1 mice. β-actin was used as the loading control. (B) Representative H&E, Masson’s trichrome, and Sirius Red staining of kidneys from the indicated groups. Scale bar, 50 μm. (C) Representative immunofluorescence images of α-SMA in kidneys from Sham + IgG, IRI-28 d + IgG, and IRI-28 d + anti-SPP1 mice. Nuclei were counterstained with DAPI. Scale bar, 50 μm. (D) Representative immunofluorescence images of SPP1 in kidneys from the indicated groups. Nuclei were counterstained with DAPI. Scale bar, 50 μm. (E–H) Quantification of Masson-positive area (E), Sirius Red-positive area (F), α-SMA-positive area (G), and SPP1-positive area (H) in the indicated groups. (I) Renal hydroxyproline content in Sham + IgG, IRI-28 d + IgG, and IRI-28 d + anti-SPP1 mice. (J) Ratio of injured kidney weight to contralateral kidney weight (IRI/CL) in the indicated groups. Data are shown as mean ± SD; indicated pairwise comparisons were performed using two-tailed unpaired Student’s t tests. **P < 0.01; ***P < 0.001.

### Post-injury Trem2 deletion suppresses fibrosis across ischemic and obstructive kidney injury

Most critically, we tested whether the upstream macrophage decoder remains required after the initiating insult. Tamoxifen-inducible Cx3cr1-CreERT2/+;Trem2fl/fl mice were compared with Cre-negative Trem2fl/fl littermates (Fig. 9A,B), with recombination induced only after IRI or UUO to minimize effects on the initial injury phase. Trem2 expression in sorted CD45 CD11b F4/80 kidney macrophages was reduced by approximately 98% in cKO mice (Fig. 9C,D), confirming efficient depletion in the renal macrophage compartment.

**Figure 9.**
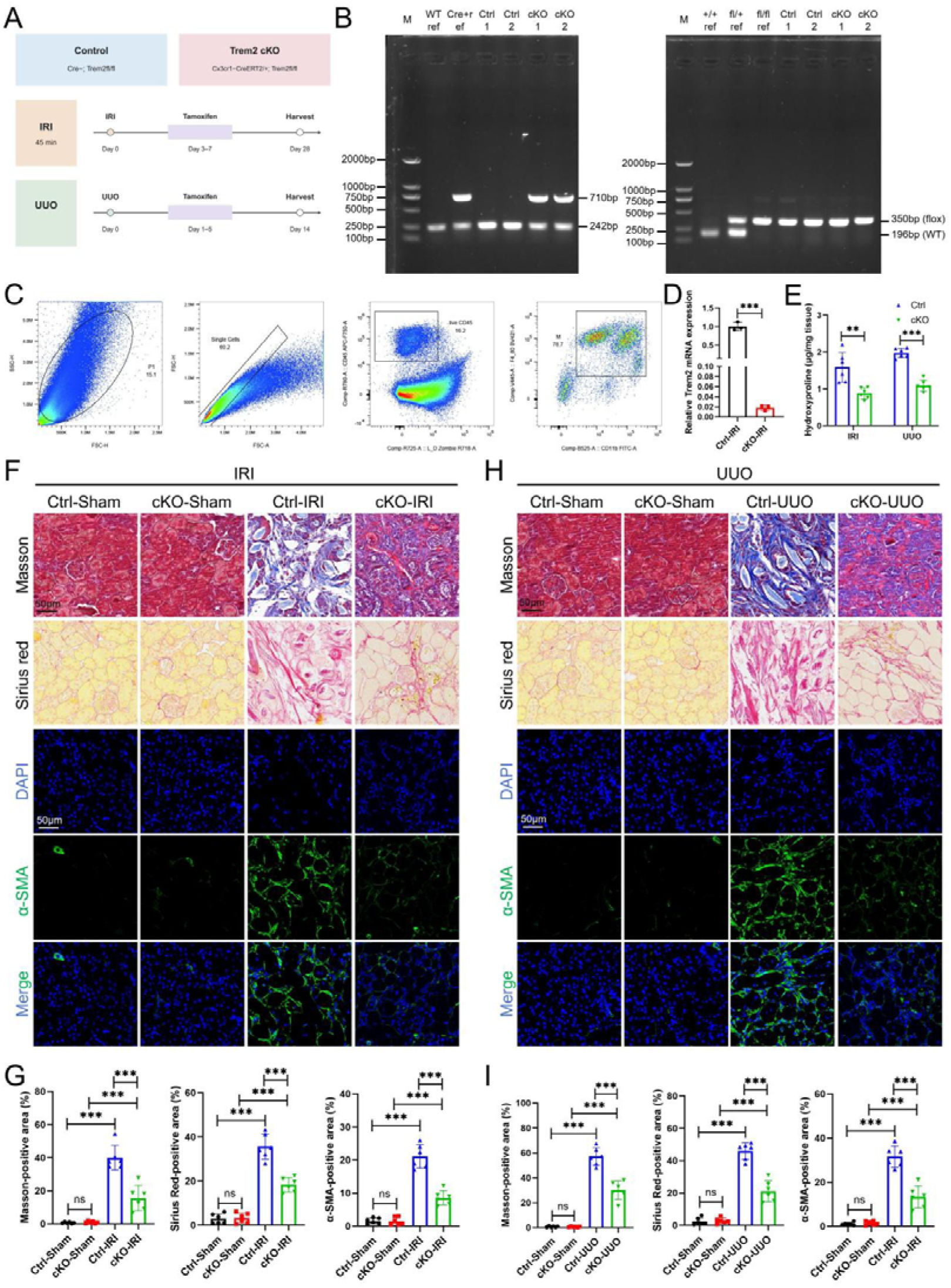
Post-injury Trem2 deletion suppresses fibrosis across ischemic and obstructive kidney injury. (A) Experimental design for inducible Trem2 deletion in Cx3cr1-CreERT2/+; Trem2fl/fl mice. Trem2fl/fl Cre-negative littermates served as controls. Unilateral renal IRI was induced on day 0, followed by tamoxifen administration on days 3–7 and tissue collection on day 28. UUO was performed on day 0, followed by tamoxifen administration on days 1–5 and tissue collection on day 14. (B) Representative genotyping PCR of the Cx3cr1-CreERT2 and Trem2 floxed alleles. For the Cx3cr1-CreERT2 assay, the wild-type and Cre-associated products migrated at approximately 242 and 710 bp, respectively. For Trem2 genotyping, the wild-type and floxed alleles migrated at approximately 196 and 350 bp, respectively. (C) Representative flow-cytometric gating strategy used for isolation of kidney macrophages. Sequential gating identified the major cellular population, singlets, viable CD45+ leukocytes, and CD11b+F4/80+ renal myeloid/macrophage cells. (D) Trem2 mRNA expression in flow-sorted kidney macrophages from Ctrl-IRI and cKO-IRI kidneys. Expression was normalized to Rplp0 and is shown relative to the Ctrl-IRI group. n = 3 mice per group. (E) Renal hydroxyproline content in Ctrl and cKO mice subjected to IRI or UUO. n = 6 mice per group. (F) Representative Masson’s trichrome staining, Sirius Red staining, and α-SMA immunofluorescence in Ctrl-Sham, cKO-Sham, Ctrl-IRI, and cKO-IRI kidneys. Nuclei were counterstained with DAPI. (G) Quantification of Masson-positive area, Sirius Red-positive area, and α-SMA-positive area in the IRI cohort. n = 6 mice per group. (H) Representative Masson’s trichrome staining, Sirius Red staining, and α-SMA immunofluorescence in Ctrl-Sham, cKO-Sham, Ctrl-UUO, and cKO-UUO kidneys. Nuclei were counterstained with DAPI. (I) Quantification of Masson-positive area, Sirius Red-positive area, and α-SMA-positive area in the UUO cohort. n = 6 mice per group. Data are shown as mean ± SD; indicated pairwise comparisons were performed using two-tailed unpaired Student’s t tests. *P < 0.05, **P < 0.01, ***P < 0.001; ns, not significant. Scale bars, 50 μm.

After IRI, cKO kidneys showed reduced Masson- and Sirius Red-positive collagen, interstitial α-SMA, and hydroxyproline versus Ctrl-IRI kidneys (Fig. 9E–G). Similar protection was observed after UUO, with reduced collagen accumulation, α-SMA expansion, and hydroxyproline in cKO kidneys (Fig. 9E,H,I). Protection across ischemic and obstructive injury indicates that repair-phase TREM2 dependence is not restricted to one injury context. Because recombination began only after injury, this experiment temporally separates TREM2-dependent fibrotic repair from effects on the initiating insult.

### Human kidney transcriptomics converge on a related TREM2/SPP1 macrophage–stromal architecture

We next asked whether the multicellular architecture defined experimentally in mice was represented in human kidney disease. In GSE211785, reclustering of the human monocyte/macrophage compartment identified a TREM2-high C4 state and a C3-like C5 state enriched for the murine C3 signature, including TREM2, SPP1, GPNMB, LGALS3, APOE, FABP5, CTSD, CTSL, ATP6V0D2, and MFGE8 (Fig. 10A–D). Of 134 TREM2 SPP1 macrophages, 99 localized to C4/C5, corresponding to 4.17% of C4/C5 macrophages versus 1.81% of other macrophages. This enrichment persisted after donor stratification (common odds ratio, 2.26; 95% CI, 1.51–3.36; P = 2.59 × 10□□; Fig. 10E–G).

**Figure 10.**
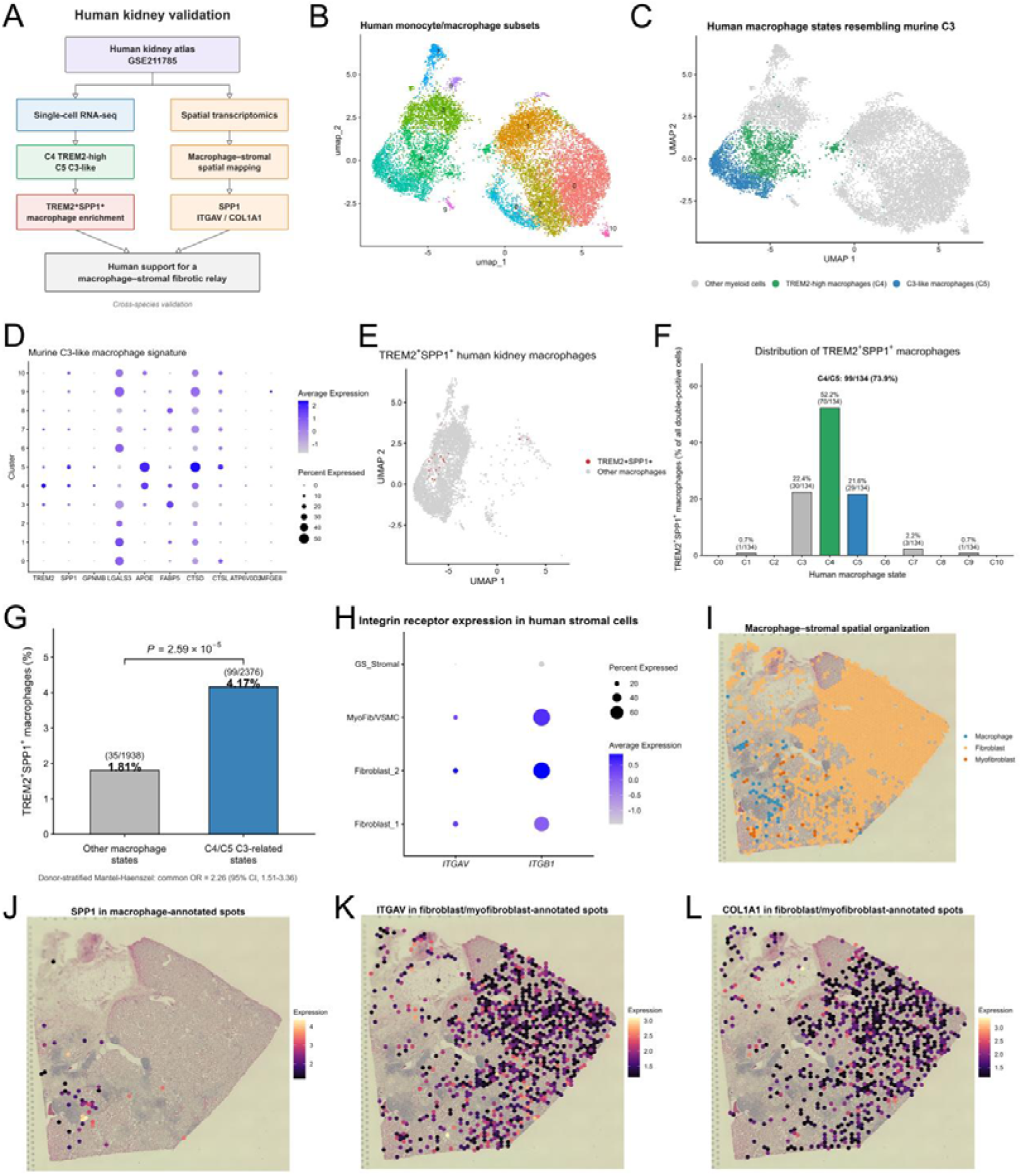
Human kidney transcriptomics converge on a related TREM2/SPP1 macrophage–stromal architecture. (A) Schematic overview of the human kidney validation strategy using the GSE211785 dataset. Single-cell RNA sequencing was used to identify human macrophage states related to the murine C3 program and to assess TREM2/SPP1 co-expression, whereas spatial transcriptomics was used to map macrophage- and stromal-associated fibrotic programs within human kidney tissue. (B) UMAP visualization of human monocyte/macrophage subsets identified in the single-cell RNA-seq dataset. (C) UMAP highlighting human macrophage states transcriptionally related to the murine C3 population, including a TREM2-high C4 population and a C3-like C5 population. Other myeloid cells are shown in gray. (D) Dot plot showing expression of the murine C3-associated macrophage signature across human macrophage clusters. Dot size denotes the proportion of expressing cells and color intensity denotes mean expression. (E) UMAP visualization of TREM2□SPP1□ human kidney macrophages identified by transcript co-detection. (F) Distribution of TREM2□SPP1□ macrophages across human macrophage states. Of 134 detected TREM2□SPP1□ macrophages, 99 (73.9%) were localized within the C4/C5 populations, including 70 cells in C4 and 29 cells in C5. (G) Frequency of TREM2□SPP1□ macrophages in C4/C5 versus the remaining macrophage states. TREM2□SPP1□ cells accounted for 4.17% (99/2376) of C4/C5 macrophages compared with 1.81% (35/1938) of macrophages in other states. Donor-stratified Mantel–Haenszel analysis yielded a common odds ratio of 2.26 (95% CI, 1.51–3.36; P = 2.59 × 10□□). (H) Dot plot showing expression of ITGAV and ITGB1 across human stromal populations, including fibroblast, myofibroblast/vascular smooth muscle cell, and stromal subsets. Dot size represents the percentage of expressing cells and color intensity indicates average expression. (I) Spatial transcriptomic map illustrating macrophage-, fibroblast-, and myofibroblast-annotated regions within human kidney tissue. (J) Spatial distribution of SPP1 expression in macrophage-annotated spots. (K) Spatial distribution of ITGAV expression in fibroblast/myofibroblast-annotated spots. (L) Spatial distribution of COL1A1 expression in fibroblast/myofibroblast-annotated spots.

Human stromal populations expressed ITGAV and ITGB1 (Fig. 10H), while spatial transcriptomics placed macrophage-associated programs within fibroblast/myofibroblast-rich regions of diseased kidney tissue (Fig. 10I–L). Although these data do not establish ligand–receptor causality in human tissue, they provide cross-species and cross-scale convergence on TREM2/SPP1-associated macrophage states within an integrin- and matrix-rich stromal environment.

### Human urinary and tissue measurements link the TREM2/SPP1 program to kidney dysfunction and fibrotic burden

Finally, we asked whether the experimentally defined macrophage program was reflected in human kidney dysfunction and remodeling. Compared with healthy controls, patients with CKD showed higher serum creatinine and blood urea nitrogen together with reduced estimated glomerular filtration rate (Fig. 11A). Urinary-cell SPP1 and TREM2 mRNA were increased in CKD, whereas CD68 was unchanged (Fig. 11B), and CKD tissue showed greater interstitial α-SMA than non-neoplastic reference tissue (Fig. 11C,D).

**Figure 11.**
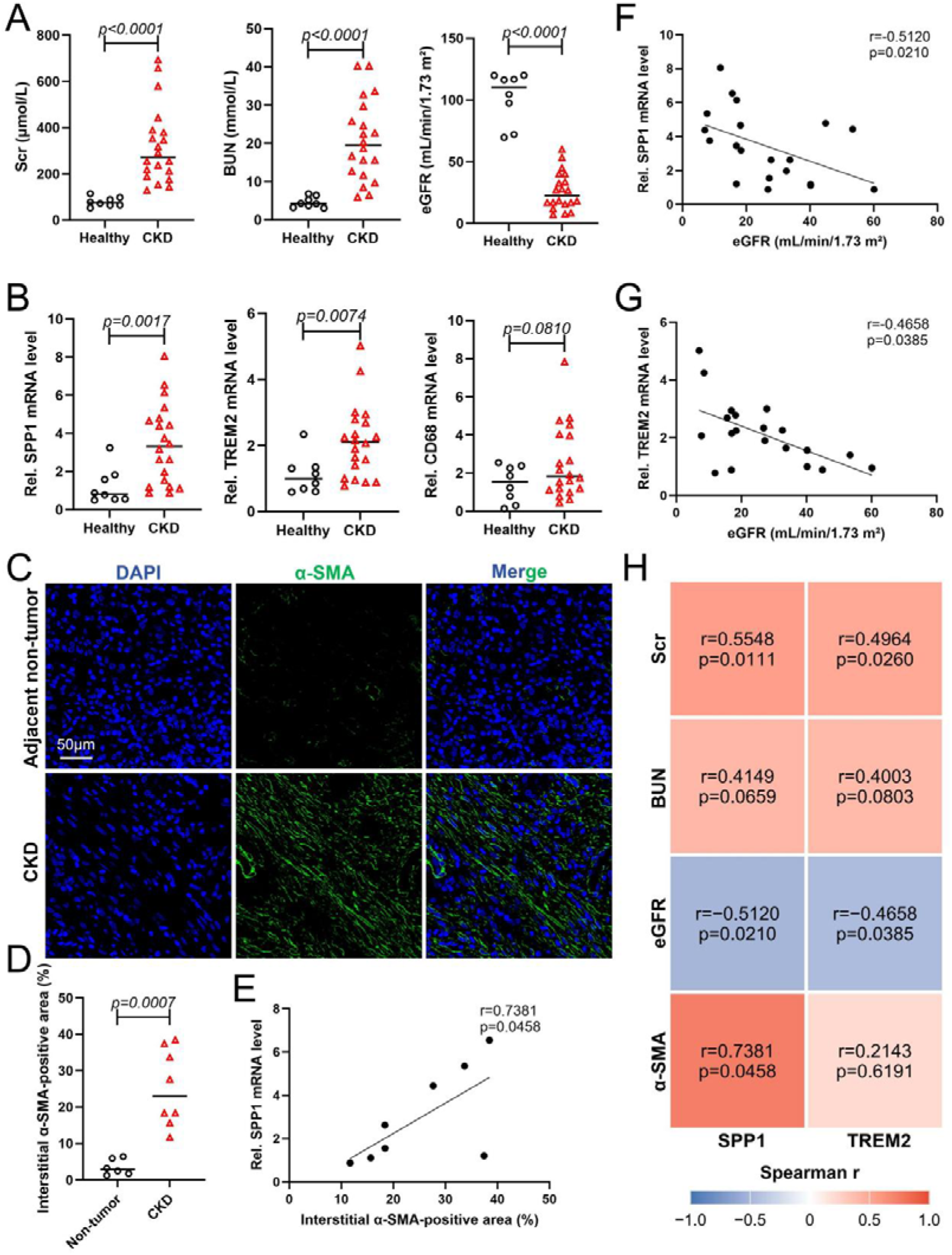
Human urinary and tissue measurements link the TREM2/SPP1 program to kidney dysfunction and fibrotic burden. (A) Serum creatinine (Scr), blood urea nitrogen (BUN), and estimated glomerular filtration rate (eGFR) in healthy controls (n = 8) and patients with CKD (n = 20). Clinical biochemical variables were compared using two-tailed Mann–Whitney U tests. (B) Relative urinary SPP1, TREM2, and CD68 mRNA levels in healthy controls and patients with CKD. Relative expression values were calculated using the 2^−ΔΔCt method and are shown for visualization, whereas statistical comparisons were performed on ΔCt values using two-tailed unpaired Student’s t tests. SPP1 and TREM2 expression was significantly increased in CKD, whereas CD68 showed no significant difference between groups. (C) Representative immunofluorescence images of α-SMA in adjacent non-tumor kidney tissues and fibrotic kidneys from patients with CKD. Nuclei were counterstained with DAPI. Scale bar, 50 μm. (D) Quantification of interstitial α-SMA-positive area in adjacent non-tumor kidney tissues (n = 6) and CKD kidney tissues (n = 8). Statistical significance was assessed using a two-tailed Mann–Whitney U test. (E) Spearman correlation between urinary SPP1 mRNA expression and interstitial α-SMA-positive area in the matched CKD tissue subset (n = 8). (F) Spearman correlation between urinary SPP1 mRNA expression and eGFR in patients with CKD (n = 20). (G) Spearman correlation between urinary TREM2 mRNA expression and eGFR in patients with CKD (n = 20). (H) Heatmap summarizing Spearman correlations of urinary SPP1 and TREM2 expression with Scr, BUN, eGFR, and interstitial α-SMA-positive area. Correlations with Scr, BUN, and eGFR were assessed in the CKD cohort (n = 20), whereas correlations with α-SMA were assessed in the matched tissue subset (n = 8). Correlation coefficients and exact P values are shown within each tile. Individual values are shown for group comparisons. All statistical tests were two-tailed.

In matched urine and tissue samples, urinary SPP1 correlated positively with interstitial α-SMA (r = 0.7381, P = 0.0458; Fig. 11E). Urinary SPP1 and TREM2 also correlated inversely with estimated glomerular filtration rate and positively with serum creatinine, whereas associations with blood urea nitrogen were not significant (Fig. 11F–H). Together with single-cell and spatial analyses, these findings link the TREM2/SPP1 program to kidney dysfunction and, for SPP1, histological fibrotic burden.

## Discussion

The principal advance of this study is the mechanistic resolution of how a transient epithelial membrane injury is converted into a persistent multicellular fibrotic state. Rather than representing parallel responses to tissue damage, tubular epithelial cells, macrophages, and fibroblasts form a directional information-transfer system in which both the cellular carrier and the physical form of the signal change across compartments. Tubular membrane injury generates a phosphatidylserine (PS)/oxidized PS (OxPS)-rich lipid context; macrophage TREM2 decodes this membrane state through separable ligand-recognition and adaptor-coupling steps; receptor signaling is transcriptionally converted into an SPP1 output; and fibroblasts receive this output through ITGAV/ITGB1-associated signaling to execute matrix remodeling. Importantly, this relay remains biologically required after the initiating injury has occurred, as demonstrated by repair-phase SPP1 neutralization and post-injury Trem2 deletion, with the latter providing protection across both ischemic and obstructive kidney injury. These findings extend the remodeling-macrophage paradigm beyond the identification of a profibrotic cell state by explaining how epithelial damage is progressively transformed across cellular compartments into a persistent fibrotic program[19]. Maladaptive repair therefore emerges not simply as the cumulative consequence of earlier injury, but as an active and interruptible process sustained by continued information transfer between injured epithelium, immune cells, and stroma.

The upstream lipid findings suggest that membrane injury itself can function as a form of biological information. H/R did not merely perturb intracellular lipid metabolism; it altered the composition and extracellular accessibility of membrane phospholipids in a manner that could be sensed by macrophages. H/R remodeled PS species, increased OxPS abundance and PS externalization, and generated tubular membrane material capable of activating TREM2–DAP12–SYK signaling. Masking exposed PS attenuated this response, whereas defined PS- and OxPS-containing liposomes reproduced key features of macrophage activation. Two orthogonal binding approaches further demonstrated direct interaction of PS/OxPS-containing lipid preparations with the TREM2 extracellular domain. Thus, the relevant upstream signal is better viewed not as a single soluble damage mediator, but as an altered membrane context through which epithelial injury becomes readable to the immune system. TREM2 is well suited to interpret such membrane environments because its macrophage functions are strongly shaped by local lipid states. In atherosclerosis, TREM2 supports lipid uptake and survival of foamy macrophages, whereas in pulmonary fibrosis local lipid ligands sustain TREM2-dependent profibrotic macrophage activity[20, 21]. Our findings add kidney-specific source information by tracing this ligand context back to injured tubular epithelium. They do not require one OxPS molecular species to act as the exclusive ligand; rather, they support PS/OxPS-containing injured membranes as a biologically active context through which damaged tubules can present injury information to macrophages.

The receptor-level experiments further show that macrophages do not passively transmit this information but actively transform it. R47A markedly impaired PS/OxPS binding while largely preserving TREM2–DAP12 association, whereas K183A disrupted DAP12 coupling despite preserved receptor abundance and membrane localization. These complementary defects separate two sequential operations within TREM2 damage sensing: extracellular phospholipid recognition and intracellular adaptor engagement. Both ultimately converged on reduced SYK activation and diminished SPP1 output. Downstream, ChIP-qPCR and promoter mutagenesis showed that NF-κB and AP-1 directly program Spp1 transcription. The macrophage therefore performs a true signal transformation rather than passive signal transmission: a membrane-associated lipid input is decoded at the receptor surface, partitioned into ligand-recognition and adaptor-coupling steps, and then rewritten as a transcriptionally programmed soluble output. This mechanistic separation also provides a framework for understanding the context-dependent biology of TREM2 in kidney disease. TREM2-dependent efferocytosis, phagocytosis, and macrophage survival may contribute to early recovery, whereas persistent engagement during unresolved repair may stabilize a fibrotic macrophage program. A disease-promoting role for macrophage TREM2 has also been reported in obstructive nephropathy, while TREM2 blockade can reshape profibrotic macrophage lipid programs in other organs[22, 23]. More broadly, our data are consistent with the idea that the duration and tissue context of damage sensing may determine whether TREM2-associated macrophage activity contributes to injury resolution or becomes incorporated into a chronic remodeling program[24].

SPP1 provides the next change in signal modality by converting macrophage-state reprogramming into stromal execution. Scar-associated macrophages during AKI-to-CKD progression display SPP1-centered communication networks, but the present experiments extend this observation beyond co-expression and inferred ligand–receptor connectivity. Conditioned-medium transfer, SPP1 neutralization, recombinant SPP1 stimulation, and genetic or antibody-mediated interference with ITGAV/ITGB1 converged on an ITGAV/ITGB1-associated FAK–SMAD2/3 response in kidney fibroblasts. These findings are consistent with SPP1-driven SMAD signaling in kidney fibrosis and with the antifibrotic effects of inhibiting αvβ1-containing integrins[25–27]. The resulting sequence represents a genuine change in signal modality across cellular compartments: epithelial membrane lipid becomes macrophage receptor signaling; receptor signaling becomes a transcriptional SPP1 output; soluble SPP1 becomes fibroblast integrin signaling; and stromal signaling becomes contractile and matrix-producing behavior. In this way, a membrane lesion that is initially confined to injured epithelium acquires persistence at the multicellular level through successive signal transformations. Repair-phase SPP1 neutralization functionally closes this downstream arm of the relay in vivo, reducing FAK–SMAD2/3 activation, matrix accumulation, hydroxyproline content, and tissue fibrosis after the initial kidney injury has already occurred.

The temporal intervention experiments are particularly important because they distinguish mechanisms that initiate injury from mechanisms that sustain maladaptive repair. Many experimental interventions are applied before or at the time of tissue injury, making it difficult to determine whether subsequent reductions in fibrosis simply reflect attenuation of the initiating insult. Here, Trem2 recombination was induced only after IRI or UUO had been established, and SPP1 neutralization likewise began during the repair phase. Despite this delay, both interventions reduced subsequent fibrosis. Moreover, post-injury Trem2 deletion was protective across two mechanistically distinct forms of kidney damage: transient ischemia–reperfusion and sustained ureteral obstruction. These findings indicate that the TREM2–SPP1 axis is not merely a molecular footprint left by earlier epithelial injury but remains functionally required as repair progresses. The concept of persistence is therefore central to the mechanism: once initiated, the macrophage–fibroblast relay continues to contribute to fibrosis even after the original tissue insult is no longer the sole determinant of disease progression. This temporal separation also carries translational relevance, because AKI is often recognized only after substantial epithelial injury has already occurred. A pathway that remains biologically active during repair represents a fundamentally different therapeutic opportunity from one whose relevance is restricted to the initiating insult.

Two additional observations support the robustness and translational relevance of this framework. First, the antifibrotic effect of delayed Trem2 deletion across both IRI and UUO argues that repair-phase TREM2 dependence is not restricted to a single mode of kidney injury. The initiating lesions are different, yet both converge on a macrophage-associated fibrotic program that remains sensitive to post-injury Trem2 deletion. Second, human data converge on a related multicellular architecture across several analytical scales. Human single-cell analysis identified TREM2/SPP1-associated macrophage states enriched for components of the murine program, while spatial transcriptomics localized macrophage-associated SPP1/GPNMB signals within integrin- and matrix-rich fibrotic regions. Urinary and kidney-tissue measurements further linked components of this program to kidney dysfunction and fibrotic burden. Human CKD studies have likewise identified macrophage–fibroblast communication within fibrotic niches, while FAK provides a mechanosensitive signaling node linking fibroblast activation to matrix production[28, 29]. Together with evidence that kidney macrophage communication is dynamically remodeled by local niche signals[30]. These observations support cross-species conservation of the broad macrophage–stromal logic. We nevertheless interpret the human data as cross-species and cross-scale convergence rather than proof of one-to-one equivalence between mouse and human macrophage states, formal biomarker performance, or causal TREM2–SPP1 signaling in human tissue.

The study also highlights a broader distinction between the presence of a damage-associated molecule and the biological persistence of damage information. PS exposure itself is transient at the level of an individual membrane, yet its recognition can be converted into a macrophage transcriptional state whose output persists beyond the original membrane event. SPP1 then transfers that information into a stromal signaling program capable of sustaining matrix production and tissue remodeling. The biological consequence is therefore not simply the sum of three independently activated cell types, but the sequential propagation of information in which each compartment changes the form of the signal it receives. This may help explain why tissue fibrosis can continue to evolve after the initiating insult has diminished: persistence does not necessarily require the original signal to remain unchanged, but may instead arise because the signal is repeatedly reformatted as it passes between cellular compartments. In the present system, membrane lipid information is converted into receptor signaling, receptor signaling into transcriptional output, and transcriptional output into stromal behavior.

Several limitations define the boundaries of this model and the most informative next experiments. First, although PS masking, defined liposomes, lipid-overlay assays, and liposome co-sedimentation provide convergent evidence for PS/OxPS–TREM2 recognition, PS/OxPS availability was not directly manipulated in vivo. The animal experiments therefore establish a repair-phase requirement for TREM2 and SPP1, whereas ligand specificity is defined primarily by cellular and biochemical studies. Annexin V masking may also alter the handling of injured membrane material and cannot by itself establish PS as the only active component of tubular debris. Second, the binding assays do not provide equilibrium affinities or identify which oxidized PS species most strongly engage TREM2. Quantitative lipid–receptor biophysics will therefore be required to define relative ligand affinities and molecular selectivity. Third, Cx3cr1-CreERT2 targets a broader mononuclear-phagocyte lineage rather than an exclusively kidney-macrophage population, although Trem2 was efficiently depleted in flow-sorted kidney macrophages. More lineage-restricted approaches would further refine the cellular requirement for TREM2 during repair. Finally, the clinical cohort was modest and designed for mechanistic validation rather than biomarker development. Larger and independently replicated human cohorts will be required to determine whether urinary or tissue components of this program have diagnostic or prognostic value. Accordingly, the present data support PS/OxPS-containing injured membranes as an upstream ligand context, while establishing TREM2 and SPP1 as repair-phase requirements in vivo and demonstrating that the macrophage–stromal relay remains interruptible after injury. Together, these findings support a model in which tubular membrane damage is progressively reformatted across cellular compartments into a persistent but interruptible fibrotic program (Fig. 12).

**Figure 12.**
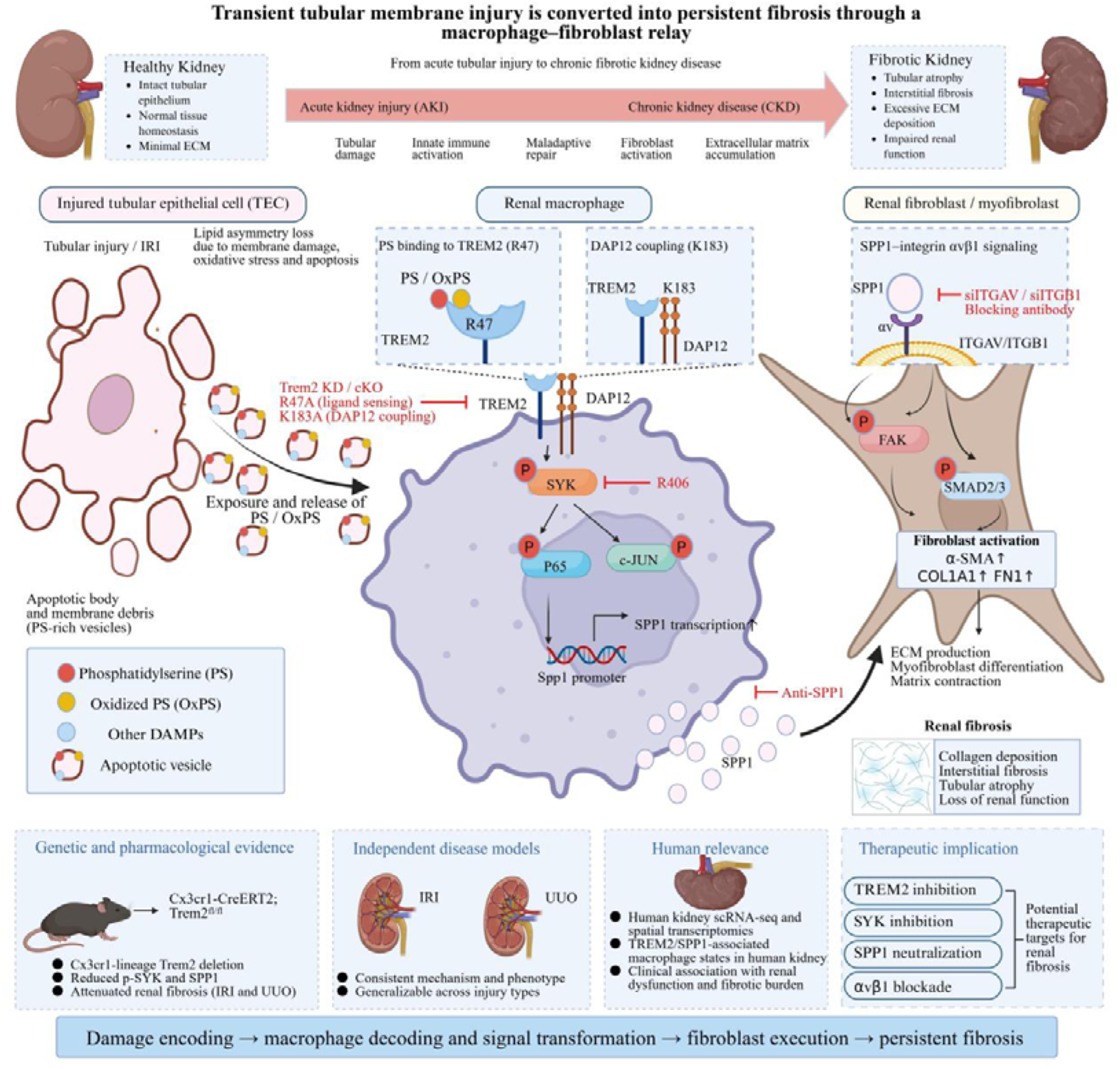
Proposed model of a tubular membrane damage–macrophage–fibroblast fibrotic relay during kidney repair. Tubular epithelial injury is accompanied by phosphatidylserine (PS) remodeling, oxidation and externalization, generating a PS/OxPS-rich membrane context that can be sensed by macrophages. TREM2 decodes this membrane-associated signal through R47-dependent phospholipid recognition and K183-dependent coupling to DAP12, leading to SYK activation and downstream NF-κB/AP-1 signaling. This receptor input is transcriptionally converted into SPP1 production, thereby transforming a membrane-lipid signal into a soluble macrophage output. Macrophage-derived SPP1 is subsequently received by kidney fibroblasts through ITGAV/ITGB1-associated signaling, activating FAK–SMAD2/3 and promoting contractile activation and extracellular-matrix production. Repair-phase SPP1 neutralization and post-injury Trem2 deletion demonstrate that this macrophage–fibroblast relay remains biologically active and interruptible after the initiating injury. The model therefore illustrates a directional sequence of epithelial damage encoding, macrophage decoding and signal transformation, and stromal execution during maladaptive kidney repair. PS/OxPS-containing injured membranes are depicted as an upstream ligand context based primarily on cellular and biochemical evidence, whereas the repair-phase requirements for TREM2 and SPP1 are supported by in vivo intervention experiments.

In summary, our findings define maladaptive kidney repair as a process of sequential information conversion across cellular compartments. Tubular membrane injury creates a phospholipid context that becomes accessible to macrophage sensing; TREM2 converts this membrane-associated input into a transcriptionally programmed SPP1 output; and fibroblasts translate that soluble signal into matrix-remodeling behavior. The persistence of this relay after the initiating insult provides a mechanistic explanation for how transient epithelial injury can acquire chronic biological consequences. Most importantly, genetic and therapeutic interventions initiated after injury remain effective, demonstrating that persistent fibrosis depends on an active signaling process rather than representing an irreversible residue of earlier tissue damage. More broadly, these findings support a framework in which the duration and transformation of damage information, rather than the initiating insult alone, help determine whether tissue repair resolves or becomes maladaptive.

## DATA AVAILABILITY

The human kidney single-cell RNA-sequencing and spatial transcriptomic datasets analyzed in this study are publicly available in the Gene Expression Omnibus under accession number GSE211785. The mouse kidney single-cell RNA-sequencing data, untargeted lipidomic data, and underlying source data generated in this study are not publicly available with this preprint. No custom code or algorithms were developed for this study.

## ACKNOWLEDGMENTS

Not applicable.

## AUTHOR CONTRIBUTIONS

Peihan Wang conceived the study, designed and performed the experiments, analyzed and interpreted the data, prepared the figures, and wrote the original manuscript. Yuan Ruan, Ting Rao and Bojun Li reviewed the manuscript. Fan Cheng and Wei Li provided funding, resources, supervision, and critical revision of the manuscript. Peihan Wang and Yuan Ruan contributed equally to this work. All authors read and approved the final manuscript.

## DECLARATION OF GENERATIVE AI AND AI-ASSISTED TECHNOLOGIES IN THE MANUSCRIPT PREPARATION PROCESS

During the preparation of this work, the authors used ChatGPT (OpenAI) to assist with language editing and improving readability. After using this tool, the authors reviewed and edited the content as needed and take full responsibility for the content of the manuscript.

